# Ancient bacterial genomes reveal a formerly unknown diversity of *Treponema pallidum* strains in early modern Europe

**DOI:** 10.1101/2020.06.09.142547

**Authors:** Kerttu Majander, Saskia Pfrengle, Judith Neukamm, Arthur Kocher, Louis du Plessis, Marta Pla-Díaz, Natasha Arora, Gülfirde Akgül, Kati Salo, Rachel Schats, Sarah Inskip, Markku Oinonen, Heiki Valk, Martin Malve, Aivar Kriiska, Päivi Onkamo, Fernando González-Candelas, Denise Kühnert, Johannes Krause, Verena J. Schuenemann

**Author notes:** these authors contributed equally (Kerttu Majander, Saskia Pfrengle).

## Abstract

Sexually transmitted (venereal) syphilis marked European history with a devastating epidemic at the end of the 15^th^ century, and is currently re-emerging globally. Together with non-venereal treponemal diseases, like bejel and yaws, found in subtropical and tropical regions, it poses a prevailing health threat worldwide. The origins and spread of treponemal diseases remain unresolved, including syphilis’ potential introduction into Europe from the Americas. Here, we present the first genetic data from archaeological human remains reflecting a previously unknown diversity of *Treponema pallidum* in historical Europe. Our study demonstrates that a variety of strains related to both venereal syphilis and yaws were already present in Northern Europe in the early modern period. We also discovered a previously unknown *T. pallidum* lineage recovered as a sister group to yaws and bejel. These findings imply a more complex pattern of geographical prevalence and etiology of early treponemal epidemics than previously understood.

## Introduction

Treponemal infections, namely yaws, bejel (endemic syphilis), and most notoriously, the sexually transmitted syphilis, represent a reoccurring, global threat to human health. Venereal syphilis, caused by *Treponema pallidum* ssp. *pallidum* (TPA) infects the worldwide human population with millions of new cases every year ^1,2^. The two endemic treponemal subspecies closely related to *T. pallidum* ssp. *pallidum* are *T. pallidum* ssp. *pertenue,* (TPE) and *T. pallidum* ssp. *endemicum* (TEN). TPE is common in the tropical regions of the world, where it causes yaws and a form of treponematosis in non-human primates. TEN is the causative agent of bejel, which is mostly found in hot and arid environments. Both of these treponematoses are usually milder in manifestations than syphilis, and have a lower incidence on the population level, yet their transmission rates have prevailed over the recent years ^3,4^. Resistance against second-line antibiotics has recently developed in *T. pallidum* ssp. *pallidum* ^5^ whereas penicillin treatment still remains effective ^1,6^. Sexually transmitted syphilis progresses slowly, while afflicting considerable damage to bone, internal organs and the nervous system ^7,8^. The endemic types of *T. pallidum* (TPE, TEN) are transmitted and mainly manifested through skin lesions, but can also affect bones and joints in a comparable way to the venereal form ^4,6^. TPA frequently transmits congenitally, resulting in various disorders for both mother and child during pregnancy, birth and infancy ^7,8^. For TPE and TEN infections today, this form of transmission is atypical, if not entirely unprecedented ^9,10^. Although the three *T. pallidum* subspecies can be separated by genetic distinctions ^11,12^, their clinical symptoms in skeletal material are difficult to distinguish ^6^. The re-emergence of syphilis is a reminder of the formidable threat it may represent with its continuous adaptations ^5,13^. Devastating outbreaks of syphilis have been documented in historical times. Early medical reports from the late 15^th^ century portray the most well-known epidemic, a rapid and Europe-wide spread of venereal syphilis in the wake of the 1495 Italian war ^14,15^. These statements also describe it having gradually changed into a milder, more chronic disease in the subsequent decades, similar in manifestation to the cases of modern day syphilis ^16,17^. These events coincide with historical expeditions, and have ignited a long-persisting hypothesis suggesting that syphilis was introduced to Europe by Columbus and his crew upon their return from the New World in 1493 ^18,19^. The alternative multiregional hypothesis contradicts this assumption, and presumes a pre-Columbian prevalence of syphilis on the European continent, potentially as a result of prehistoric spread of the disease through African and Asian routes ^20–22^. Skeletal evidence from human remains carrying pathological marks characteristic for treponematoses and dated prior to Columbus’ return from the New World, have been reported ^23,24^. However, to date no genetic evidence exists that could confirm the existence of pre-Columbian syphilis in Europe ^18,25^. Potential syphilis infections are mentioned in the literature since Medieval times, but these diagnoses may have been confounded by symptomatic similarities with other diseases, and delusively called ‘venereal’ or ‘hereditary’ leprosy ^17,25^. Misdiagnoses occurred until recently, due to the challenges in recognizing *T. pallidum* infections from other diseases and its subspecies from each other ^26^. In the present-day, the treponematoses are largely segregated in medical diagnostics either with traditional serological tests or with the more recently developed multi-locus sequence typing (MLST) schemes ^27–30^. Before the modern genetic classifications were introduced, supporters of the ‘unitarian hypothesis’ claimed that all treponematoses were in fact one and the same disease ^31,32^. Although this theory was justified in questioning the geographical distribution and clinical symptoms as the main means of categorization, the understanding of phylogenetic cladality between the treponemal subspecies has since disputed its more general principle. Genomic studies have found that the TPA and TPE strains, although clearly separated phylogenetically, remain extremely similar, and their geographical origin and time of emergence have proved complicated to confirm ^11,15,33,34^. A recent genetic study on modern lineages of treponematoses supported a common ancestor of all current TPA strains in the 1700’s ^34^, whereas the more general diversification of *T. pallidum* into subspecies has been addressed in previous studies and assumed to have happened in prehistoric times ^19,35^. Since modern genomes reflect the evolutionary situation of their isolation time, the mutation rate estimates drawn from them can be biased by natural selection yet to happen ^36^. Past lineages may also reveal lost variation unrepresented by the currently known pathogen strains available for research ^37^. For these reasons, reconstructed ancient bacterial genomes have an unprecedented potential to illuminate their species’ unresolved divergence times and origin. Several historically significant pathogens have been successfully reassembled for investigation and their reconstructed genomes have greatly contributed to our understanding of the evolution and spread of re-emerging infectious diseases ^38–40^. Ancient DNA (aDNA) studies concerning treponematoses have so far remained scarce for both biological and methodological reasons. The treponemal spirochetes survive poorly outside their host organism and are present in extremely low quantities during late stage infections, often evading detection even in living patients ^41^. The final, tertiary stage produces the most notable alterations to the skeleton in response to the human immune system, making these treponemal cases most frequently recognized, but less likely to yield genetic evidence, due to the clearance or latency of the pathogen ^42,43^. Most notably, the bones that likely contain a large amount of treponemal agents belong to congenitally infected neonates. These fragile remains rarely survive and are, even when present, often overlooked in the archaeological record ^44,45^. Previously, it would not have been feasible to use samples with low bacterial loads to detect pathogen DNA; however, recent advances in target-enrichment, high-throughput sequencing and sensitive screening methods for aDNA have aided overcoming this issue ^46^. Currently, the technical advancements, together with careful selection of samples affected by treponemal pathogenesis, are enabling genomic studies on this elusive pathogen for the first time. The means to recover *T. pallidum* from historic human remains were recently established in a study on perinatal and infant individuals from colonial Mexico, in which two ancient genomes of *T. pallidum* ssp. *pallidum* and one ancient *T. pallidum* ssp. *pertenue* genome were described^45^. However, attempts to retrieve *T. pallidum* DNA from historical adult individuals have so far been unsuccessful.

Here, we analyze ancient bacterial genomes from four novel historical *T. pallidum* strains retrieved with target enrichment from pathological human remains, including adult and subadult individuals, originating in central and northern Europe. The newly reconstructed ancient genomes represent a variety of *T. pallidum* subspecies including a formerly unknown form of treponematosis phylogenetically basal to both bejel and yaws lineages. For the first time, treponemal genomes dated temporally close the New World contact have been retrieved from European samples, including closely related strains to the endemic types today mostly restricted to the tropics and subtropics.

## Results

### Geographical origins and osteological analyses of samples

For this study, remains from nine individuals were included: five from the Crypt of the Holy Spirit in Turku, Finland, one from the Dome churchyard in Porvoo, Finland, one each from St. Jacob’s cemetery and from St. George’s cemetery in Tartu, Estonia, and finally, one from Gertrude’s Infirmary in Kampen, the Netherlands (Supplementary Table 1. a). Four of the samples from these individuals proved positive for treponemal infection in the DNA screening (Figure 1). The sample CHS119 (Figure 1.1) is a premolar from an adult individual from Turku that showed several bone changes on the skull and both tibias, indicative of treponemal infection ^47^. The sample PD28 from Porvoo (Figure 1.2) showed no lesions directly associated with syphilis ^48^ and infection was implicated primarily by the individual’s perinatal death. The sample SJ219 from a young adult from Tartu (Figure 1.3) showed putative indicators of syphilis, although the lesions present did not provide a clear distinction from mycobacterial infections like leprosy. From the young adult individual from Kampen, only a disarticulated tibia unassociated with a grave was available ^19^, which was used to obtain sample KM14-7 (Figure 1.4) directly from an active lesion on the surface. For sampling criteria, see Supplementary Note 1.

**Figure 1.**
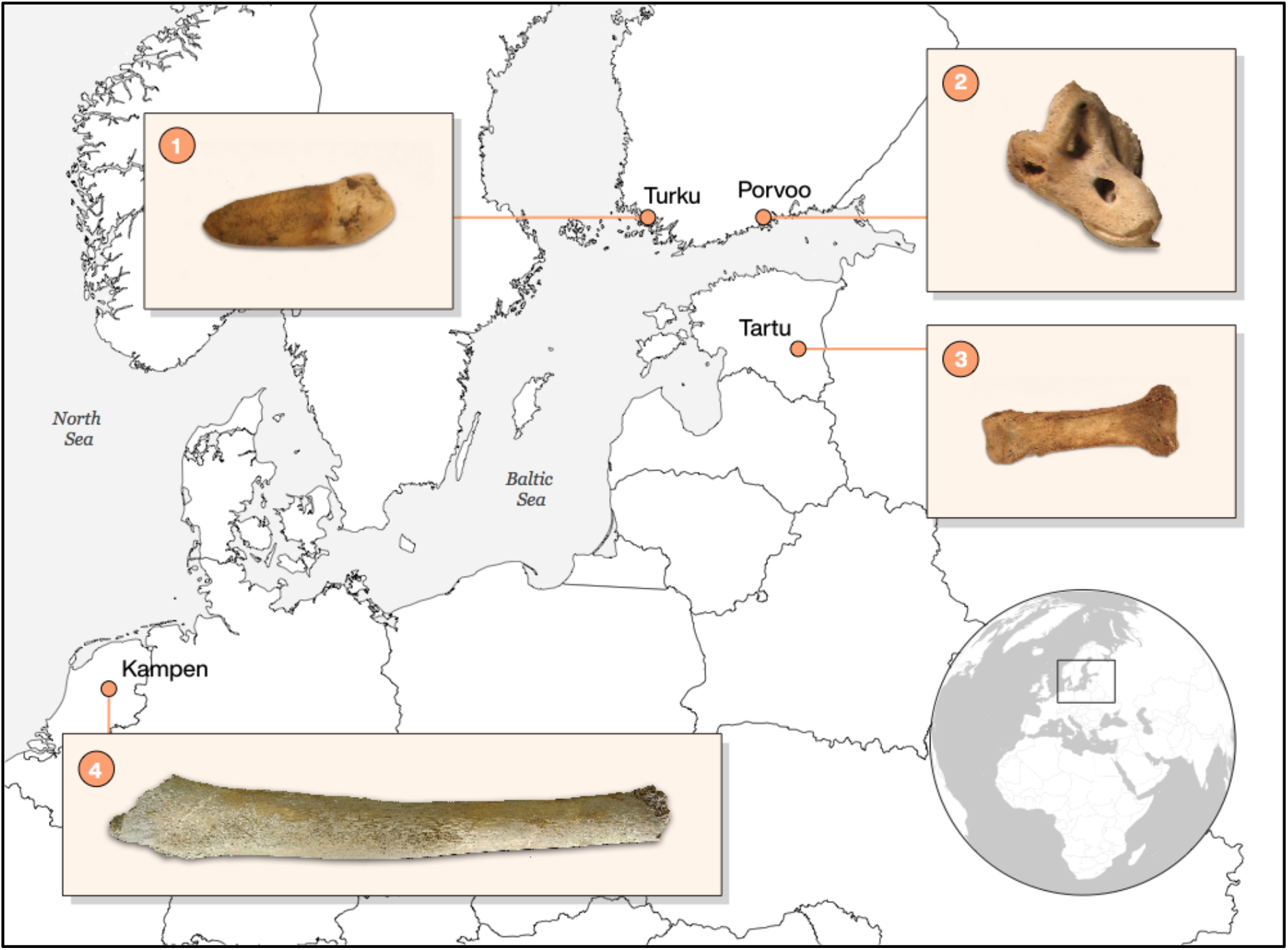
Locations of archeological sites, from which the samples used in this study originate: 1) A premolar from an adult individual 119, Crypt of the Holy Spirit, Turku, Finland, 2) petrous bone of a perinatal individual from the grave 28 at the Porvoo Dome cemetery, Porvoo, Finland, 3) a phalanx of a young adult individual from St. Jacob’s cemetery, Tartu, Estonia and 4) a tibia from a young adult individual from the Gertrude’s Infirmary, Kampen, the Netherlands.

**Figure 2:**
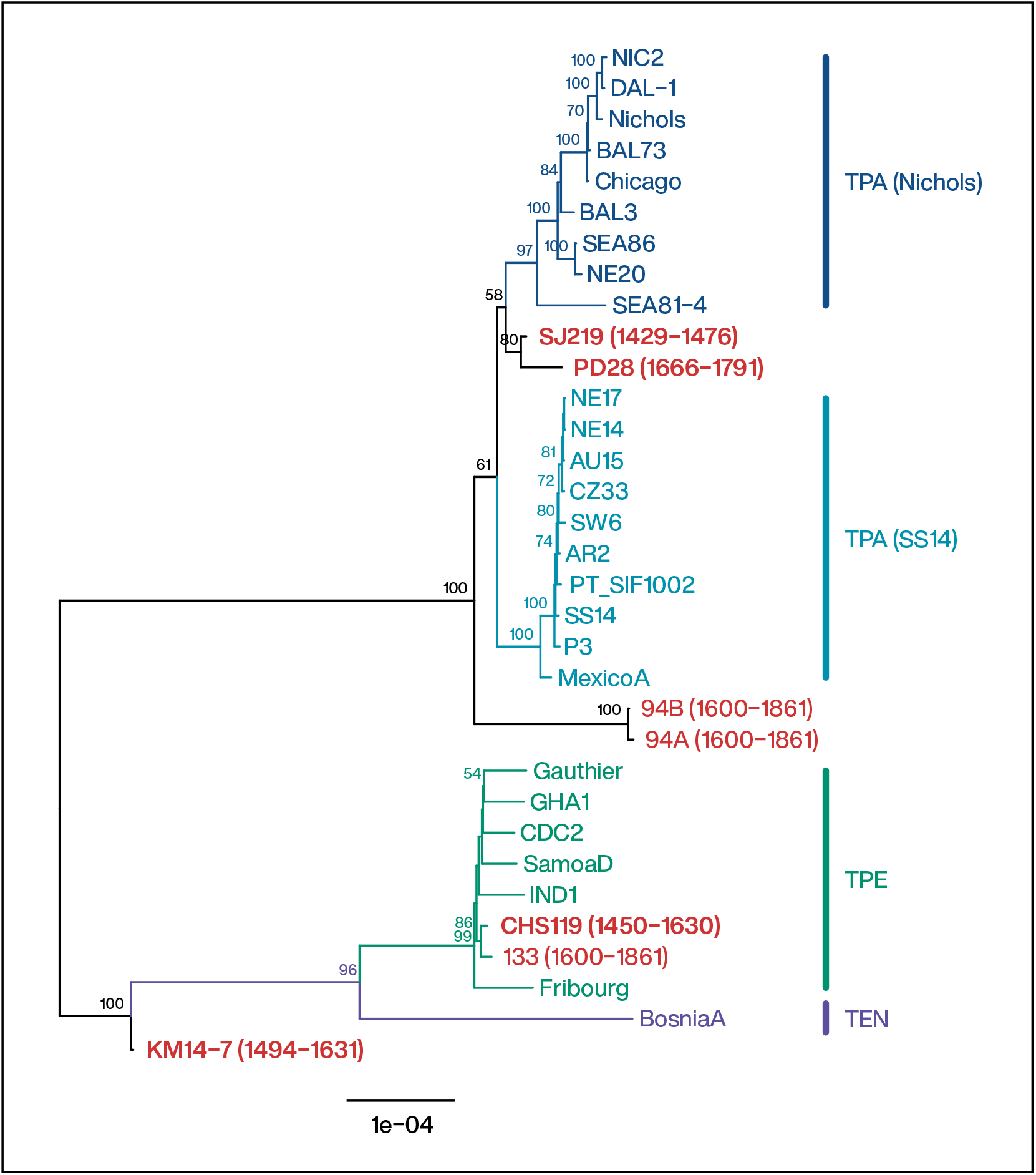
Midpoint-rooted phylogenetic tree of *Treponema pallidum* strains estimated with maximum likelihood. Branch lengths represent numbers of substitutions per site. Bootstrap support values are written at the nodes when > 50. Ancient genomes are marked in red and the ones generated in this study are in bold text.

All four positive samples were radiocarbon dated (in Klaus-Tschira-Archäometrie-Zentrum am Curt-Engelhorn-Zentrum, Mannheim, Germany, in the Laboratory of Chronology, Finnish Natural History Museum, Helsinki, and in the AMS laboratory, ETH Zürich). For PD28, the calibrated dates range from 1666 to 1950 CE. The church cemetery, however, was replaced in 1789 CE, indicating the last possible burial time for the individual ^49^. CHS119 and SJ219 both show AMS results dating the samples starting from the 15^th^ century CE. For CHS119 the upper limit of dating ranges to the 17^th^ century, whereas for SJ219, two independent laboratory estimates confirmed an upper limit within the 15th century CE. The disarticulate bone KM14-7 is C14 dated to a range from the late 15^th^ to early 17^th^ century CE^50^. Marine and freshwater reservoir effects can cause an offset in C^14^ ages between contemporaneous remains of humans or animals mainly relying on terrestrial food sources, and those principally using food sources from aquatic environments^51^. Calibration corrections for the affected radiocarbon results are feasible, but require a local baseline of isotopic signature which is currently unavailable for many regions in the world. Alternatively, other available carbon-based materials from the grave can be used to confirm the dating of an individual in uncertain cases. The putatively pre-Columbian samples CHS119 and SJ219 went through additional attestment of the dating procedures: A fragment of the wooden coffin of the individual SJ219 was used for additional dating analysis and the reservoir effect corrections were produced for the sample CHS119 ^52,53^. The resulting estimates, however, gave both individuals a date range upper limit reaching 17^th^ century CE. For more details on radiocarbon dating and reservoir effect correction, see Supplementary Note 2.

### Genome reconstruction and authenticity estimation of ancient DNA

A screening procedure using MALT ^54^ and Megan extension for visualizing ^55^ was utilized to determine *T. pallidum* positive samples. The extracts were immortalized in both double-stranded and single-stranded DNA libraries ^56–59^. Enrichment for *T. pallidum* DNA was done for all samples and in addition human mitochondrial DNA was enriched for samples CHS119, SJ219, and KM14-7 ^60,61^. After high-throughput sequencing on Illumina HiSeq4000 and NextSeq500 platforms, the resulting 98-256 million raw reads were merged sample-wise and duplicate reads removed. Genomes for each sample were then reconstructed by mapping to the Nichols reference genome (NC_021490.2) using the EAGER-pipeline ^62^. The samples yielded between 18034 and 1430292 endogenous unique reads mapping to *T. pallidum* reference and covered 47-98% of the genome at least once (Table 1. and Supplementary Table 1.b and 1.c). A total of 2637 single nucleotide polymorphisms (SNPs) were called, out of which 91 were likely spurious and were thus removed in the SNP evaluation procedure. In this procedure, a considerable number of variable positions originally identified in ancient genomes were excluded (Supplementary Table 3).

**Table 1.** Mapping statistics from the EAGER pipeline for the four historical samples screened as positive for treponemal infection. The table includes final number of mapped reads after duplicate removal, average coverage, percent of the genome covered at 2-fold coverage, percent of reads with DNA damage for the 1st base at the 5’ end and 3’ end (non-UDG treated), and average fragment length (in basepairs) for the treponemal and *T. pallidum* capture data. Same statistics are given for the human mitochondrial capture data where applicable, with the addition of estimated amount of contamination from Schmutzi program ^121^ and haplogroup assignments from HaploGrep2 ^65^ program. Molecular sexing of individuals is based on the human endogenous reads from shotgun data ^122,123^.

| Individual ID (this study) | Molecular sex | Data type | Mapped reads (post duplicate removal) | Average coverage | Genome coverage 2-fold | DNA damage 1st base 5' (non-UDG) | DNA damage 1st base 3' (non-UDG) | Average fragment length | Contamination | Haplogroup |
| --- | --- | --- | --- | --- | --- | --- | --- | --- | --- | --- |
| PD28 | XY | Treponemal | 1430292.00 | 136.23 | 98.07 | 0.04 | 0.02 | 59.91 | NA | NA |
|  |  | Mitochondrial | NA | NA | NA | NA | NA | NA | NA | NA |
| SJ219 | XX | Treponemal | 29198.00 | 1.41 | 34.00 | 0.10 | 0.11 | 49.12 | NA | NA |
|  |  | Mitochondrial | 29031.00 | 118.91 | 100.00 | 0.13 | 0.11 | 67.74 | 0.02 | HV16 |
| CHS119 | XX | Treponemal | 52054.00 | 2.82 | 62.31 | 0.18 | 0.18 | 54.16 | NA | NA |
|  |  | Mitochondrial | 4490.00 | 16.25 | 99.99 | 0.14 | 0.15 | 59.85 | 0.02 | J1c2c1 |
| KM14-7 | XY | Treponemal | 18034.00 | 0.91 | 19.35 | 0.18 | 0.17 | 51.69 | NA | NA |
|  |  | Mitochondrial | 3414.00 | 11.02 | 99.37 | 0.16 | 0.18 | 53.35 | 0.01 | U2e1f1 |

The average fragment length of the four samples varied between 49 and 60 bases, and the percentage of deaminated bases at the ends of reads, signaling authenticity of ancient DNA ^63,64^, ranged from 2-18% (*T. pallidum* DNA) and 10-18% (mtDNA). Human mitochondrial haplogroups further distinguished the geographical origin of the samples ^65,66^. The human mitochondrial DNA ranged from 3414 to 29031 mapped reads, with an average coverage of 11-119% on the mitochondrial genome. Haplogroups identified were J1c2c1 for CHS119, HV16 for SJ219, and U2e1f1 for KM14-7; all representative of the early modern variation present within northern and central European populations.

### Phylogenetic analysis and genetic recombination

Midpoint-rooted ML trees of our historical European genomes, previously published colonial Mexican genomes ^45^ and 26 modern treponemal genomes ^34,67–75^ were constructed for the alignment. The four ancient European genomes are placed at three distinct positions in the phylogenetic tree. The PD28 and SJ219 genomes most closely resemble the strains in the syphilis-causing cluster. There is considerable uncertainty regarding the exact position among the major TPA clades. However, we obtain the highest support for a position basal to the Nichols cluster (see Supplementary Figures 2 and 3). The CHS119 genome, conversely, is consistently placed in the TPE cluster of the treponemal family tree and forms a cherry with the ancient TPE genome 133 from colonial Mexico. The affinity of these two ancient genomes may in part result from similarly missing data with respect to the complete genomes of the modern strains in the clade. The central European KM14-7, remarkably, falls basal to all TPE lineages known today, as well as to the only TEN genome published to date ^73^. This unexpected position was further supported by investigating genomic positions for which the ancestral variant of TPE/TEN and TPA clades was likely different. Among 30 of these positions for which the variant was recovered in KM14-17, 60% were TPE/TEN-like and 40% were TPA-like. In comparison, all other genomes had 100% variant characteristics of one or the other clade at these positions, except for 94A, 94B and SEA81-4 which had 16, 14.3 and 10.7% of TPE/TEN like variants respectively (Supplementary Table 4).

A recombination analysis was conducted as described in Arora *et al.* 2016^34^, including 26 modern genomes ^34,67–75^ and six ancient ones: the colonial Mexican genomes 94A, 94B, and 133 from an earlier study ^45^ and three of our historical European genomes, namely PD28, CHS119 and SJ219 (Table 2). Sample KM14-7 was excluded from the recombination analysis due to its sporadic placement in the Maximum-Likelihood (ML) tree topologies, which were derived for entire genomes and for each gene individually. Congruence between the complete genome alignments and gene trees was tested after evaluating the corresponding phylogenetic signal for each gene. For 40 loci, the phylogenetic signal and incongruence was significant. For those cases, we further verified the presence of at least three consecutive SNPs supporting a recombination event. Twelve loci passed this test and also correspond to those found as recombinant loci in a more extensive study of modern *T. pallidum* genomes (n=75) ^76^. Two of the recombining genes identified in Arora *et al*. ^22^ were also confirmed in association with the ancient European genomes. Of our ancient genomes, PD28 was possibly involved in one recombinant event of the TP0136 gene as a putative recipient, along with the Nichols clade, with the TPE/TEN clade, CHS119 and colonial Mexican 133 genomes as putative donors. The same possibility was observed in the recombination event detected in the TP0179 gene, although only with TPE/TEN clade and 133 as presumptive donors. One putative recombination event concerning the TP0865 gene was identified between the TPE/TEN clade, including the CHS119 and 133 genomes and the SEA86, NE20 and SEA81-4 lineages. Finally, there is another recombination event concerning the TP0558 gene, with the TPE/TEN clade and CHS119 genome as potential donors and the SS14 clade, MexicoA, 94A and 914B from colonial Mexico, and PD28 genomes as recipients. Other assumed events involving recombination events between the modern strains and the previously published ancient genomes from the New World are listed in Supplementary Table 5.

**Table 2.** Genes involved in putative recombination events involving the historical European TPA genome PD28 and TPE genome CHS119. Slashes are used to separate the different potential gene donor strains. The affected recipient strains are separated by commas. Arrows point to the likely direction of recombination between the donor and recipient strains. An interrogation mark indicates and uncertain yet likely involvement in the event.

| Gene ID | Event | Start | End | Minimal size | Strains involved |
| --- | --- | --- | --- | --- | --- |
| TPANIC_0136 | 5 | 973 | 1034 | 61 | Yaws/Bosnia/133/CHS119? -> PD28, Nichols clade |
| TPANIC_0179 | 1 | 259 | 647 | 388 | Yaws/Bosnia/133 -> PD28, Nichols clade |
| TPANIC_0558 | 1 | 414 | 834 | 420 | Yaws/Bosnia/CHS119?/133? -> PD28, 94B, 94A?, SS14 clade, MexicoA |
| TPANIC_0865 | 1 | 258 | 576 | 318 | Yaws/Bosnia/CHS119?/133? -> Sea86, Ne20, Sea81-4 |

### Molecular clock dating

Molecular clock dating analyses were performed on a 28 genome dataset, described in detail in the Methods section Molecular clock dating. Linear regression of root-to-tip genetic distance against sampling date indicated that the *T. pallidum* strains possess a moderate temporal signal (*r* = 0.467) (Figure 4). The significance of the correlation was confirmed using a permutation test (*p* = 0.015). The correlation is higher within the TPA and TPE clades alone (r = 0.742 and r = 0.832, respectively). Within the TPA cluster the signal is driven by the SS14 clade (r = 0.852), while the Nichols clade does not appear clock-like (r = −0.007).

The Bayesian date randomization test (DRT) showed that neither the mean nor the median estimate of the molecular clock rate estimated under the true sampling dates was contained within the HPD intervals of estimates from any of the replicates with permuted sampling dates (Supplementary Figure 4). In addition, the HPD intervals of the clock rate est Figure imates of only 3 of the 50 replicates intersect with the HPD interval estimated under the true sampling dates. We interpret these results as evidence that the molecular clock signal in the dataset is substantially stronger than expected by chance. Finally, robustness analyses conducted using different combinations of demographic and clock models showed that the demographic model has little effect on the estimated divergence times and sampling dates and that a relaxed clock model receives strong support (see Supplementary Note 3).

Posterior distributions of divergence dates of *T. pallidum* clades and sampling dates of historical genomes are shown in Figure 4a and Figure 4b shows the MCC tree estimated in BEAST v.2.6. Median estimates and 95% HPD intervals are given in Supplementary Tables 6.B and 6.C. T_MRCA_ (time for the most recent common ancestor) calculated for the whole *T. pallidum* family is placed far in the prehistoric era, at least 2500 BCE. However, time-dependency of molecular rates (TDMR) may lead to underestimating deep divergence times when mutation rates are inferred from genomes collected within a relatively restricted time period ^77,78^. Applying a model accounting for TMDR may be possible, but would require an inclusion of genomes sampled over wide and distinct time-periods ^79,80^. The latest common ancestor of the venereal syphilis strains was placed between the 10^th^ and 15^th^ century CE. The divergence of TPE and TEN (yaws and bejel) was dated between the 9^th^ century BCE and the 10^th^ century CE, while the most recent common ancestor of TPE was placed between the 14^th^ and 16^th^ century CE. Among the TPA strains the T_MRCA_ of the Nichols clade (13^th^ to 17^th^ century CE) was clearly older than that of the SS14 clade (18^th^ to 20^th^ century CE). Due to the inclusion of 4 historical genomes, the above divergence times are substantially older than the times reported in Arora et al (2016) ^34^. Similarly, the estimated mean molecular clock rate (median estimate 1.037 × 10^−7^ s/s/y, 95% HPD 6.856 × 10^−8^ – 1.447 × 10^−7^ s/s/y) is slower than the clock rates reported in recent studies ^5,34^ for either *T. pallidum* as a whole or for TPA strains exclusively. Nonetheless, the 95% HPD of the mean clock rate overlaps with the estimates previously reported 5. For more information, see Supplementary Figure 5).

Molecular clock dating allows us to refine the sampling date estimates of three of the four historical genomes (Figure 4b). The posterior distributions of the sampling dates of PD28 and CHS119 place most of the weight on more recent dates, while that of 133 favours an older sampling date. This is especially pronounced for CHS119 and 133 with the 95% HPD interval not including any dates older than 1526 CE for CHS119 or younger than 1773 CE for 133. On the other hand, for SJ219, the 95% HPD of the sampling date spans nearly the entire range defined by radiocarbon dating, making it impossible to exclude a pre-Columbian sampling date (posterior probability 0.26) (Supplementary Table 6. C).

### Virulence factor analysis

A virulence factor analysis for the historical European strains was conducted as first introduced by Valtuena *et al.* in 2017 ^81^, on a set of 60 TPA genes associated with virulence functions outlined in previous studies ^34,45,82,83^. The sequenced reads were mapped with quality threshold as 0, and annotated according to the Nichols reference genome (NC_021490.2). The observed coverage of four modern TPA lineages, including the SS14 lineage and three Nichols clade strains NIC2, Sea81-4 and Chicago ^34,67,74^, compared with the four historical treponemal genomes from Europe and three treponemal genomes from colonial Mexico ^45^ are visualized in a heatmap (Figure 5), showing an evenly distributed coverage for putative virulence genes for the European ancient genomes. The PD28 genome shows a full and CHS119 a relatively high coverage, whereas for the samples SJ219 and KM14-7 the coverage is generally lower. All virulence factors, including *Tpr* genes, which exist in all of the *Treponema pallidum* subspecies and code for porin structures in the cell membrane ^12,84,85^ are entirely covered in the novel ancient European genomes, as well as the *FadL* gene found missing in the colonial Mexican TPA genomes 94A and 94B ^45^. More in-depth nucleotide level comparisons between the historical genomes and modern strains are needed to further investigate the differences in the protein coding functions related to these genes in the past.

## Discussion

### Early emergence of syphilis in Europe

In this study, four ancient *Treponema* genomes were retrieved from human skeletal remains dating to early modern Europe, providing unprecedented insights into the first reported epidemics of syphilis at the end of the 15^th^ century. Two of our ancient genomes, PD28 and SJ219, were identified as TPA strains, the causative agent of syphilis, representing the first molecularly identified specimen of *T. p. pallidum* from historical Europe. These genomes fall within the modern variety of the TPA strains. They form a sister clade to the modern TPA branch relatively basal to all lineages, although their precise position regarding the two major clades, Nichols and SS14, could not be recovered with high confidence. The TPA-carrying sample PD28 was placed in the early modern period by combined analyses of archaeological context and C^14^ dating. Two independent radiocarbon analyses were performed on the sample SJ219 from an Estonian Medieval archaeological context, both indicating that the individual dated from early to mid-15^th^ century, distinctly before Columbus’ expeditions. This dating would point to the TPA contagions occurring in Europe prior to the New World contact, suggesting that the causative agent of the continent-wide epidemic at the end of the 15^th^ century may have resided within Europe before contact. However, a reservoir effect could have influenced the radiocarbon analysis results of the individual SJ219 through her diet, and the dating could not be confirmed with complete certainty (see Supplementary Note 1 and Supplementary Table 1.a. for more information). Such direct evidence of TPA from an early European context gives unprecedented support for the existing prevalence of venereally transmitted syphilis around and potentially prior to the contact Columbus initiated in the Americas.

### Yaws in Europe

Of our four historical genomes, two fall outside the variation of TPA. One of them, the genome CHS119 from Finland, clusters with the present TPE group (causative agent of yaws). Although the direct radiocarbon dating places the sample in the 15^th^-16^th^ century, a full confidence of the exact date cannot be gained, due to the potential marine reservoir effect ^52,53^. This sample provides the first evidence of yaws infections in historical Northern Europe, far from the tropical environment in which present-day yaws is typically found. Strikingly, the contemporaneous genome KM14-7 from the Netherlands falls basal to both the bejel-causing lineage and all known strains causing yaws, unveiling a previously unidentified lineage of *T. pallidum*.

Due to the endogenous DNA coverage retrieved for the KM14-7 sample, the inclusion of this genome in the recombination and time-calibrated phylogenetic analysis was limited. Despite this, the ML tree topology with the KM14-7 in the basal position to TPE and TEN clades could be further confirmed with a closer nucleotide level inspection. The lineage shows genetic similarities to both currently existing syphilis (13 unique SNPs) and yaws (25 unique SNPs), but it represented a distinct form from both, and had apparently diverged from their common root before the cluster consisting of yaws and bejel today, that we dated to at least 1000 years BP. Altogether, these different ancient treponemal genomes from northern and central Europe point to an early existing variety of *T. pallidum* in the Old World. Their existence does not refute the potential introduction of new strains of treponemes from the New World in the wake of the European expeditions, but provides credibility to an endemic origin of the 15^th^ century epidemics.

Whereas recombination events between the three modern day treponemal subspecies are deemed rare ^76,86^, such events were observed across the subspecies in our study. These recombination events presumably happened in the past, before the geographical niches were acquired by the TPA, TPE, and TEN agents ^34,45,76^. The historical cases of syphilis and yaws in an overlapping area provides a plausible opportunity for recombination. The potential recombination events observed in this study involved both the lineages present in the modern day variation and the ancient genomes PD28 and CHS119 from Europe and 94A and 914B genomes from Mexico ^45^.

Overall, our observations point towards recombination events happening in the direction of syphilis-causing clades from the yaws and bejel-causing ancestors. These recombinations between the clades further support a geographically close common history of the TPA and TPE lineages, which cannot be concluded from the geographical distribution of modern day lineages.

### Old hypotheses revisited

The reconstructed genomes from historical Finnish and Estonian human remains show an early spread of both venereal syphilis and yaws at the northern end of Europe. The diversity and geographical extent of these *Treponema* lineages recovered close to the contact period suggest the presence of treponematoses in the Old World prior to the first recorded epidemic events. Although the C^14^ analyses and the archaeological context of the individuals CHS119 and SJ219 support the pre-Columbian dating, these claims are thwarted by the uncertainties in the dating methodology (see Supplementary Note 2). In addition to the direct dating of the individual samples, we used our novel historical genomes for molecular dating of the phylogenetic clades of *T. pallidum*. The dating analysis sets the T_MRCA_ for the entire *T. pallidum* family to less than 2000-3000 BCE. The available calibration points, however, provide only the lower bound for the divergence of the different subspecies and permit the hypotheses according to which Treponemas may possess a deep prehistoric background in association to their human hosts ^20,21^. The T_MRCA_ for all TPA clade strains between 10^th^ and 15^th^ century CE supports a radiation of these strains within Europe instead of having a single ancestral source from the New World. The branching of the yaws and bejel clade between 9^th^ century BCE and 10^th^ century CE clearly predates contact period, and together with the genome KM14-7 with ancestral characteristics, suggests a common history of these diseases in the Old World. All the currently known lineages of yaws have a common ancestor as late as 14^th^ to 16^th^ century CE, which could point to a novel radiation simultaneously with the rise of the early venereal syphilis, possibly enabled by the same evolutionary opportunities around the contact period, or due to the competition between the concurrent subspecies.

The basal position of the ancient genome KM14-7 in the phylogenetic tree of *T. pallidum* represents a possibly unprecedented type of treponemal infectious agent. The lineage it belonged to carried genetic similarities to both currently existing syphilis and yaws, yet it appears to be a distinct form from both. The pathogenesis of this agent may have resembled the endemic types of treponematoses, since the majority of its recovered SNPs are shared with the yaws and bejel lineages. It has been suggested that yaws or its ancestors represent the original form of treponematosis that appeared and spread around the world thousands of years ago, was re-introduced to the Iberian Peninsula via the Central and Western African slave trade, some 50 years before Columbus’ travels, and eventually gave rise to the venereal syphilis ^35,87^. It is indeed possible that the more severe venereal form in the Old World developed from a mild endemic type of disease, enhanced by genetic recombination events or in response to a competition between the various existing pathogens ^31,84^. Likewise, recombination events may have occurred between the endemic European strains and novel lineages introduced at the wake of the New World contact, precipitating the epidemic events at that time. While cladality between the different subspecies clearly exists in both the past and modern day, it now seems likely that recombination has interconnected these clades in the past, and that the genetic differences do not necessarily define the treponemal pathogenesis observed in the archaeological remains. Since diagnostic signs in skeletal remains are hard to distinguish between yaws and venereal syphilis, and so far undiscovered early treponematoses may have existed simultaneously, only further genetic studies on samples originating from all continents can properly address hypotheses about the direction of spread and order of the epidemic events. Presumably many past treponemal lineages remain unknown today and, once revealed, will prove pivotal in uncovering the relationship between treponemal strains and in dating their emergence.

### Outlook and implications on sampling strategies

Retrieving treponeme’s DNA from skeletal material is highly challenging, and the feasibility of the effort has been strictly questioned before the recently published colonial Mexican genomes ^42,43,45^. Here, four out of the nine included individuals yielded a sufficient amount of treponemal aDNA for in-depth genomic analyses (Supplementary Table 1. a). While the previously published Mexican genomes were obtained from neonates and infants only ^45^, we were able to recover *T. pallidum* DNA also from subadult (KM14-7, SJ219) and adult (CHS119) individuals, including one with only a tentative diagnosis of the disease (SJ219). Using bone tissue directly involved with ongoing inflammation or possessing an ample blood flow, such as an active lesion (KM14-7) and dental pulp (CHS119), probably facilitated the successful sampling, although in the case of the neonate (PD28), even a petrous bone proved highly successful, yielding an entire genome of historical TPA strain up to 136-fold. This first retrieval of pathogen DNA from a petrous bone was likely due to a systemic condition related to an early congenital infection with an extremely high bacterial load ^88^.

Notably, one of the colonial Mexican infants from a previous study ^45^ likely suffered from yaws infection, as well as one of the Finnish individuals (CHS119) in this study. These cases lend credibility to the notion that the different treponemal agents cause essentially similar skeletal alterations, and are highly adaptable to environmental circumstances ^33,89^. We therefore propose that the geographical separation criteria between the treponemal diseases should be used with caution, especially when it comes to earlier forms of treponematoses and their diagnostic manifestations in the archaeological record. Overall, the reconstruction of novel treponemal genomes from these various ancient sources further proves the feasibility of retrieving treponemal aDNA from skeletal material and raises the hope of achieving progress with the prevailing cases of advanced and latent infections. Improving methodologies targeted for samples with low bacterial load and genomic coverage may soon aid in recovering positive aDNA results from putative cases of treponematoses from early-to pre-historic contexts, thereby illuminating the most persistent quandaries of the field, such as the ultimate origin of venereal syphilis.

## Materials and Methods

### Sample processing

Documentation and UV-irradiation of the bone material for decontamination, as well as laboratory procedures for sampling, DNA extraction, library preparation, and library indexing were all conducted in facilities dedicated to ancient DNA work at the University of Tübingen, with necessary precautions taken including protective clothing and minimum contamination-risk working methods.

All post-amplification steps were performed in a laboratory provided by the Department of Toxicology at the University of Tübingen as well as in the post-PCR laboratory of the Paleogenetics Group, Institute of Evolutionary Medicine (IEM), University of Zurich (UZH). DNA sequencing was performed at the sequencing facilities of the Max Planck Institute for the Science of Human History in Jena or at the Functional Genomics Center at the University of Zurich (FGCZ).

### Sampling and DNA extraction

Before extracting DNA from the samples, all surfaces were irradiated with ultraviolet light (UV-irradiated) to minimize potential contamination from modern DNA. DNA extraction was performed according to a well-established extraction protocol for ancient DNA ^56^. For DNA extraction, 30-120 mg of bone powder was used per sample. The bone powder was obtained by drilling bone tissue using a dental drill and dental drill bits. For different individuals, variable amounts of extracts were produced. During each extraction one positive control (ancient cave bear bone powder sample) and one negative control were included for every ten samples. Positive controls for extractions were carried along until the indexing of DNA-libraries, and the negative controls were carried through all following experiments and sequenced. Sample-specific barcodes (indexes) were added to both ends of the libraries ^57^. The indexed libraries were then amplified to reach a DNA concentration of 90ng/μl, at a minimum. The amplification was performed using 1x Herculase II buffer, 0.4 μM IS5 and 0.4 μM IS6 primer ^57^, Herculase II Fusion DNA Polymerase (Agilent Technologies), 0.25 mM dNTPs (100 mM; 25 mM each dNTP) and 5 μl indexed library, as template. Four reactions per library were prepared for a total amplification reaction volume of 100μl. The thermal profile included an initial denaturation of 2 minutes at 95 °C, then 3-18 cycles depending on the DNA concentration after indexing of the libraries, followed by another denaturation for 30 seconds at 95 °C, 30-second annealing at 60 °C, and 30-second elongation at 72 °C.

### Library preparation

In this study, double-stranded (ds) and single-stranded (ss) DNA-libraries were produced. All DNA library preparation procedures that were applied in this study are described in the following paragraphs.

#### Double-stranded DNA library preparation

For the preparation of DNA-libraries 20μl of DNA extract was converted into ds-DNA libraries ^58^. For every ten libraries produced, one negative control was used. Negative controls for library preparations were carried along in parallel with all following experiments. Sample-specific barcodes (indexes) were added to both ends of the libraries ^57^. The indexed libraries were then amplified to reach a minimum DNA concentration of 90ng/μl. The amplification was performed using 1x Herculase II buffer, 0.4 μM IS5 and 0.4 μM IS6 primer^57^, Herculase II Fusion DNA Polymerase (Agilent Technologies), 0.25 mM dNTPs (100 mM; 25 mM each dNTP) and 5 μl indexed library as DNA template. Four reactions per library were prepared and the total amplification reaction volume was 100μl. The thermal profile included an initial denaturation for 2 minutes at 95 °C and 3-18 cycles, depending on DNA concentration after indexing of the libraries, denaturation for 30 seconds at 95 °C, 30 seconds annealing at 60 °C and a 30-second elongation at 72 °C, followed by a final elongation step for 5 minutes at 72 °C. All splits of one indexed library were pooled and purified using the Qiagen MinElute PCR purification kit. The final quantification for all DNA libraries was performed with the Agilent 2100 Bioanalyzer.

#### Uracil-DNA Glycosylase (UDG) treated double-stranded DNA library preparation

To avoid potential sequencing artifacts caused by the characteristic ancient DNA damage profile ^90^ additional libraries for genome-wide enrichment were created, namely 30μl of DNA extract was pre-treated with uracil-DNA glycosylase ^91^. Controls were also treated accordingly. Sequencing libraries were indexed and amplified as described above.

#### Single-stranded DNA library preparation

Single-stranded libraries were generated from 20 μl of DNA extract according to established protocols ^59^. Two single-stranded libraries were prepared for each of four individuals. All single-stranded libraries were indexed and amplified with the same experimental procedures as applied and described for the double-stranded DNA libraries. To assess the library concentration, D1000 High Sensitivity ScreenTape was used on an Agilent 2200 Tapestation. All libraries were pooled in equimolar concentration for the genome-wide enrichment step.

### Capture techniques

#### Capture for a first screening for treponemal DNA

For the first screening for *Treponema pallidum* DNA, all double-stranded libraries without UDG treatment for all individuals were pooled. All negative controls were pooled separately. The *T. pallidum* screening was performed using a modified version ^34^ of the array capture approach originally developed by Hodges and colleagues ^60^ to enrich the sample libraries for *T. pallidum*-specific DNA. Here, we used arrays designed by Agilent Technology and the blocking oligonucleotides (BO) 4, 6, 8, and 10 ^57^ in the initial screening and followed the same enrichment procedure as described by Arora and colleagues ^34^ and Schuenemann and colleagues ^45^, as follows:

#### Whole-genome capture

After the analysis of the shotgun screening data from the initial sequencing, UDG-treated double-stranded DNA libraries and single-stranded DNA libraries were produced for samples potentially positive for syphilis and used for whole genome capture. The whole-genome capture was performed as described above using the same array enrichment strategy. In addition to the blocking oligonucleotides for double-stranded libraries, specific blocking oligonucleotides 4, 6, 8, and 11 ^57^ were used for single-stranded libraries. The whole-genome enrichment for treponemal DNA was produced in three rounds of array capture and a maximum of two libraries from different individuals were pooled for each array. Enrichment pools were diluted to 10 nMol/L for sequencing.

### In-solution capture for KM14-7

An additional in-solution capture procedure was performed for sample KM14-7 to obtain higher coverage. Genome-wide enrichment of single-stranded libraries was performed through custom target enrichment kits (Arbor Bioscience). RNA baits with a length of 60 nucleotides and a 4bp tiling density were designed based on three reference genomes (Nichols, GenBank: CP004010.2, SS14: GenBank CP000805.1, Fribourg Blanc: GenBank CP003902). Five hundred ng library pools were enriched according to the manufacturer’s instructions. Captured libraries were amplified in 3×100 μl reactions containing 1 unit Herculase II Fusion DNA polymerase (Agilent), 1x Herculase II reaction buffer, 0.25mM dNTPs, 0.4 μM primers IS5 and IS6 ^57^ and 15 μl library template. The thermal profile was the following: initial denaturation at 95**°**C for 2 min, 14 cycles of denaturation at 95**°**C for 30 sec, annealing at 60**°**C for 30 sec, and elongation at 72**°**C for 30 sec, followed by a final elongation at 72**°**C for 5 min. Captured libraries were purified with MinElute spin columns (QIAGEN) and quantified with a D1000 High Sensitivity ScreenTape on an Agilent 2200 TapeStation.

### Sequencing

The first two rounds of genome-wide enriched (array enrichment strategy) double-stranded libraries were sequenced at the Max Planck Institute for the Science of Human History in Jena on an Illumina HiSeq 4000 platform with 1*75+8+8 cycles (single-end reads), following the manufacturer’s protocols for multiplex sequencing. The third round of genome-wide array capture enrichment for *Treponema pallidum* DNA by double-stranded libraries was sequenced at the Max Planck Institute for the Science of Human History in Jena on an Illumina HiSeq 4000 platform with 2*50+8+8 cycles (paired-end reads), following manufacturer’s protocols. In the fourth round of sequencing the single-stranded DNA libraries, enriched by array capture strategy for *Treponema pallidum* DNA, were sequenced. The sequencing was performed on an Illumina NextSeq500 platform 2*75+8+8 cycles (paired-end reads) at the Functional Genomics Center Zurich.

### Read Processing, Mapping and Variant Calling

The capture data from the sequencing runs were merged sample-wise and data processing was performed using EAGER version 1.92.37 (Efficient Ancient GEnome Reconstruction) ^62^. The quality of the sequencing data was assessed using the FastQC ^92^ and the reads were adapter trimmed with AdapterRemoval ver. 2.2.1a ^93^. All samples were mapped to the Nichols reference genome (NC_021490.2), using CircularMapper version 1.0^62^ with a minimum quality score of 37 and a stringency parameter n=0.1. Duplicates were removed with MarkDuplicates v2.15.0 ^94^. MapDamage version 2.0.6 was used to investigate the damage patterns, which can authenticate the ancient origin of the DNA sequences (Supplementary Figure 1). The Genome Analysis Toolkit (GATK) version 3.8.0 ^95,96^ was used to perform SNP calling and generate vcf files for each genome.

### Genomic dataset and multisequence alignment

We constituted a genomic dataset representative of the extant diversity of *T. pallidum* and including the three previously published ancient genomes from Mexico (Supplementary Table 2). Raw sequencing data was gathered for strains that were high-throughput sequenced. The previously exposed procedure was then applied to obtain vcf files for each genome. We then used MultiVCFAnalyzer ^97^ to produce alignments with the following parameters: bases were called if covered by at least two reads with a mapping quality of 30 and a consensus of at least 90% (with the one-read-exception rule implemented in MultiVCFAnalyzer). The resulting alignment was realigned with already assembled genomes (isolates BosniaA, CDC2, Chicago, Fribourg, Gauthier, MexicoA, PT_SIF1002, SS14, and SEA81_4_1), using AliView version 1.21 ^98^.

### SNP quality assessment

Calling SNPs from ancient bacterial DNA data is challenging due to DNA damage, potential environmental contamination and low genome coverage which may lead to the recovery artifactual genetic variation in reconstructed DNA sequences. This can interfere with all subsequent analyses and, in particular, lead to artificially long branches in phylogenetic trees and impede time-calibrated analyses. Artifactual SNPs resulting from environmental contamination shared between several samples may also lead to biases in inferred phylogenetic tree topologies or generate misleading evidence of genetic recombination.

In order to filter artifactual SNPs, we used the SNPEvaluation tool ^99^ as proposed by Keller and colleagues ^100^. More specifically, for all newly generated ancient genomes, as well as for all previously published genomes for which the mean sequencing coverage was below 20, we reviewed any unique SNP and any SNP shared by less than 6 genomes that had at least one of the following features in a 50-bp window around the SNP: (i) some positions were not covered, (ii) the reference was supported by at least one read or (iii) the coverage changed depending on the mapping stringency (i.e. we compared the initial alignments with “low-stringency” alignments produced with bwa parameter n=0.01). Any SNP supported by less than four reads was excluded (i.e. N was called at that position) if at least one read supported the reference or if the SNP was “damage-like” (i.e resulting from a C to T or G to A substitution). Furthermore, the specificity of the reads supporting the SNPs was verified by mapping them against the full GenBank database with BLAST ^101^. Any SNP supported by reads mapping equally or better to other organisms than *T. pallidum* was excluded. Since many of these false SNPs likely arising from non-specific mapping were located in tRNAs, we excluded all tRNAs from the alignments. The list of excluded positions was written in a gff file (Supplementary Data 1) and removed from the full alignment generated by MultiVCFAnalyzer still contains excluded SNPs, which we removed using an in-house bash script.

### Phylogenetic and recombination analysis

An analysis pipeline described by Pla-Díaz and colleagues ^76^ was used to investigate putative recombining genes, which could potentially interfere with phylogenetic tree topologies between the modern and ancient lineages. Altogether 12 putative recombinant genes were subsequently excluded, resulting in an alternative positioning of the ancient TPA strains close to the Nichols cluster. The pipeline included the following steps: 1) Obtaining an ML tree with the complete multiple alignment. 2) Obtaining ML tree for each protein-coding gene. 3) Testing the phylogenetic signal in each gene alignment. 4) Testing the congruence between each gene tree and the complete genome alignment and between each gene alignment and the complete genome tree. 5) Confirming a minimum of three consecutive SNPs congruent with a recombination event for all genes passing the filtering of step 3 and 4. 6) Removing the protein-coding genes with signals of being involved in recombination events.

After removing the recombinant genes, a phylogenetic analysis using RAxML v. 8.2.12 was performed with the rapid bootstrap algorithm based on a SNP alignment of the genomic dataset in which positions having > 25% missing data or identified as recombining were excluded. The ASC_GTRGAMMA substitution model was used together with the “stamatakis” ascertainment bias correction to account for the number of constant sites not appearing in the SNP alignment. Using the full alignment or different level of missing-data trimming had little effect on the reconstructed tree (Supplementary Figures 2-3). Not excluding identified recombining loci resulted in longer branch lengths and the placement of PD28 and SJ219 basal to the TPA SS14 clades rather than basal to the Nichols clade (Supplementary Figure 3. a).

### KM14-7 SNP analysis

In the phylogenetic tree, KM14-7 was placed basal to TPE and TEN clades. Although bootstrap support was very high, we decided to further evaluate the authenticity of this remarkable position because this genome contained a large fraction of missing data. We investigated genomic positions for which the ancestral variant of TPE/TEN and TPA was likely different. Our rationale was that if KM14-7’s position on the branch connecting TPE/TEN and TPA was authentic, the genome should contain a significant number of both TPE/TEN and TPA-like variants. In practice, we looked at positions (i) resolved in KM14-7, (ii) for which the majority variant was differing between TPE/TEN and TPA clades, but (iii) shared by more than 90% of the (modern) genomes within each clade. We then looked at the proportion of TPE/TEN and TPA-like variants in KM14-7 and compared that with all other genomes. Because KM14-7 was not included in the recombination analysis, we did not trim the recombining region for this analysis in order to avoid a bias. Finally, we also produced ML trees based on positions resolved in KM14-7 (141 SNPs). The resulting trees corresponded to the previously observed topology, with the KM14-7 recovered basal to TPE/TEN (Supplementary Figures 2 and 3).

### Molecular clock dating

KM14-7 and the Mexican TPA genomes, 94A and 94B, were removed from the molecular clock dating analyses due to their low coverage and the amount of unique SNPs present on these lineages. Furthermore, the Nichols and NIC2 genomes were removed to avoid any bias caused by continuous passaging in rabbit hosts from their isolation in the clinical context in 1912 until at least 1981 ^102–105^.

The strength of the molecular clock signal in the dataset was investigated by regressing the root-to-tip genetic distance (measured in substitutions/site) of genomes against their sampling dates ^106,107^ (Figure 3.A). Root-to-tip genetic distances were calculated on a midpoint-rooted ML tree estimated in RAxML v. 8.2.11 ^108^ using the same procedure described above (Figure 3.B). Sampling dates of historical sequences were fixed to the middle of the date range defined by radiocarbon dating (Supplementary Table 6.A). To assess the significance of the correlation we permuted the sampling dates across genomes and used the Pearson correlation coefficient as a test statistic ^109,106^(Figure 3.C). We performed 1,000 replicates and calculated the *p*-value as the proportion of replicates with a correlation coefficient greater than or equal to the truth (using the unpermuted sampling dates).

**Figure 3:**
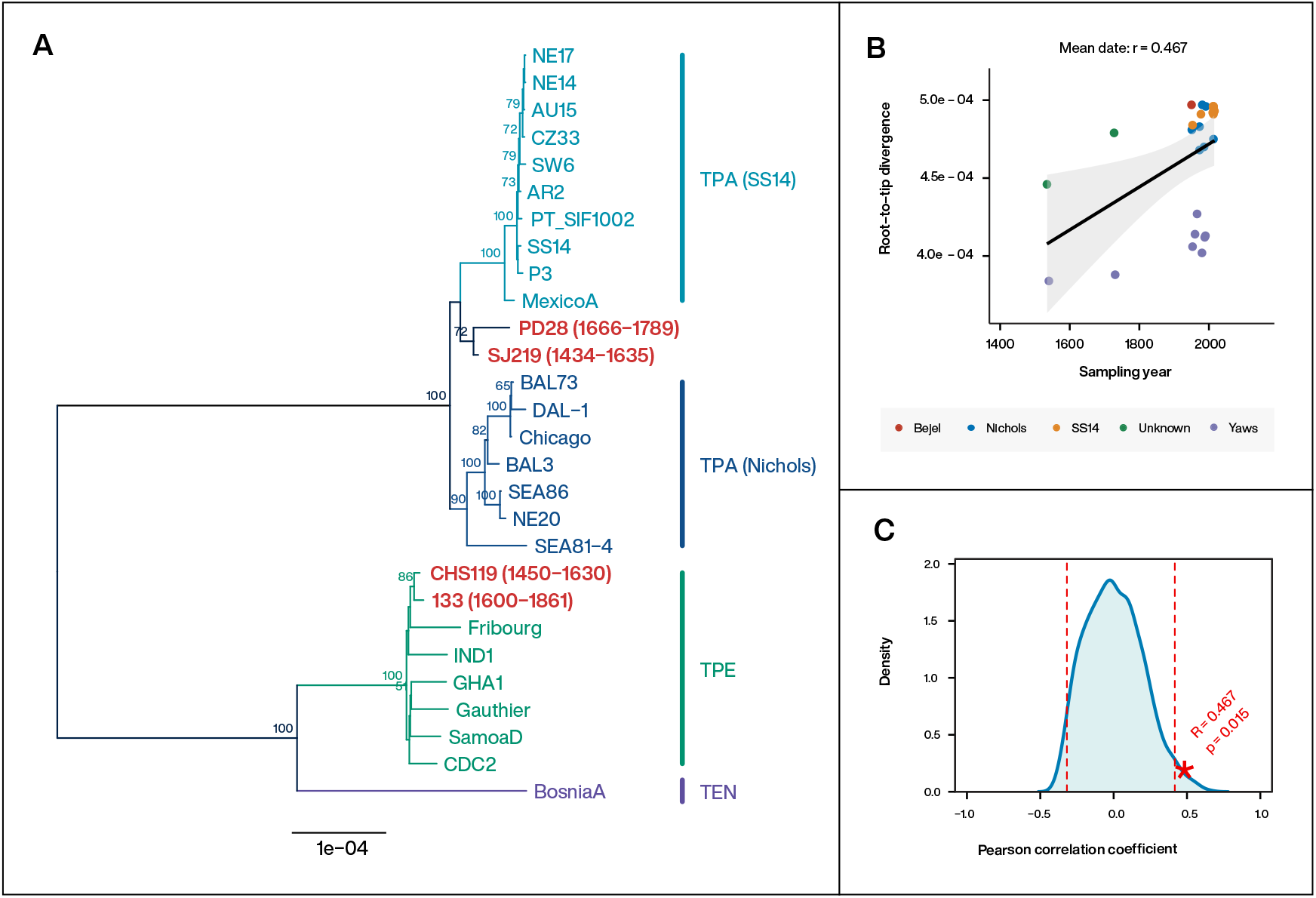
**(A)** Midpoint-rooted ML tree used to calculate root-to-tip distances in the 28 genome dataset used for molecular clock dating. The sampling date range is given in parentheses after the sample name. The historical samples from this study and the previously published colonial Mexican genome 133 are highlighted with red. **(B)** Root-to-tip genetic distance in the ML tree plotted against sampling date, using the mean date of the date range defined by radiocarbon dating. The Pearson correlation coefficient between root-to-tip distance and sampling date is given above the figure. **(C)** The null distribution of the Pearson correlation coefficient between root-to-tip distance and sampling date when permuting sampling dates across genomes (1,000 replicates). The dashed lines indicate 2.5 and 97.5 quantiles of the distribution and the star indicates the correlation coefficient using the true (unpermuted) sampling dates. The *p*-value is the proportion of replicates with the test statistic (Pearson correlation coefficient) greater than or equal to the true value.

**Figure 4:**
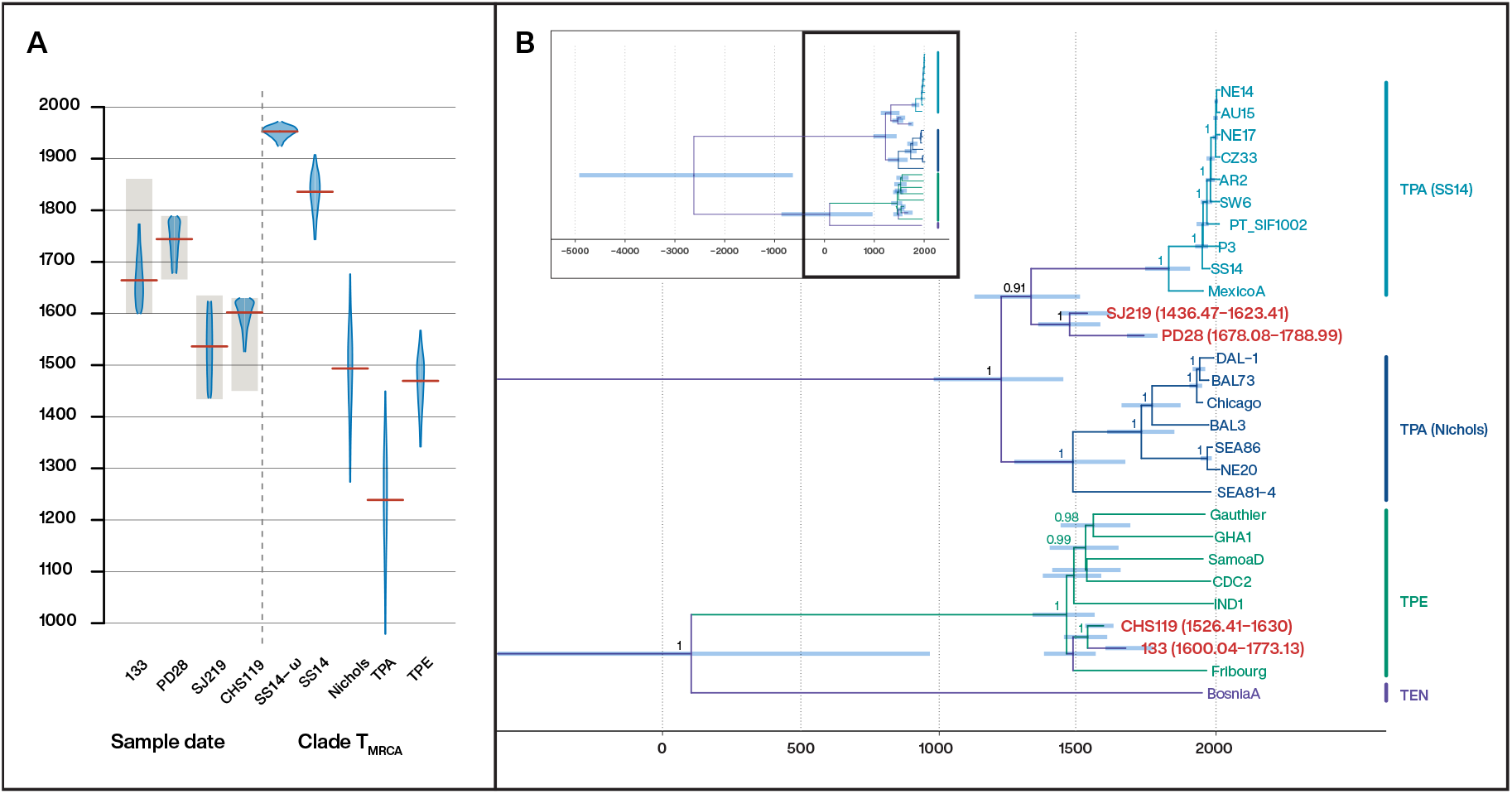
A) Posterior distributions for the sampling dates of historical genomes (left) and the divergence dates of more recent clades in the tree (right). The distributions are truncated at the upper and lower limits of the 95% HPD interval and the red lines indicate the median estimates. The shading indicates the prior distributions used for the sampling dates of historical samples (uniform priors defined by the radiocarbon date ranges). B) Maximum clade credibility (MCC) tree of the 28-genome dataset estimated in BEAST v2.6 under a relaxed clock model and a Bayesian skyline plot demographic model. Node bars indicate the 95% HPD interval of internal nodes and sampling dates of historical genomes (genomes highlighted in red and with HPD interval of the sampling dates in parentheses).

**Figure 5:**
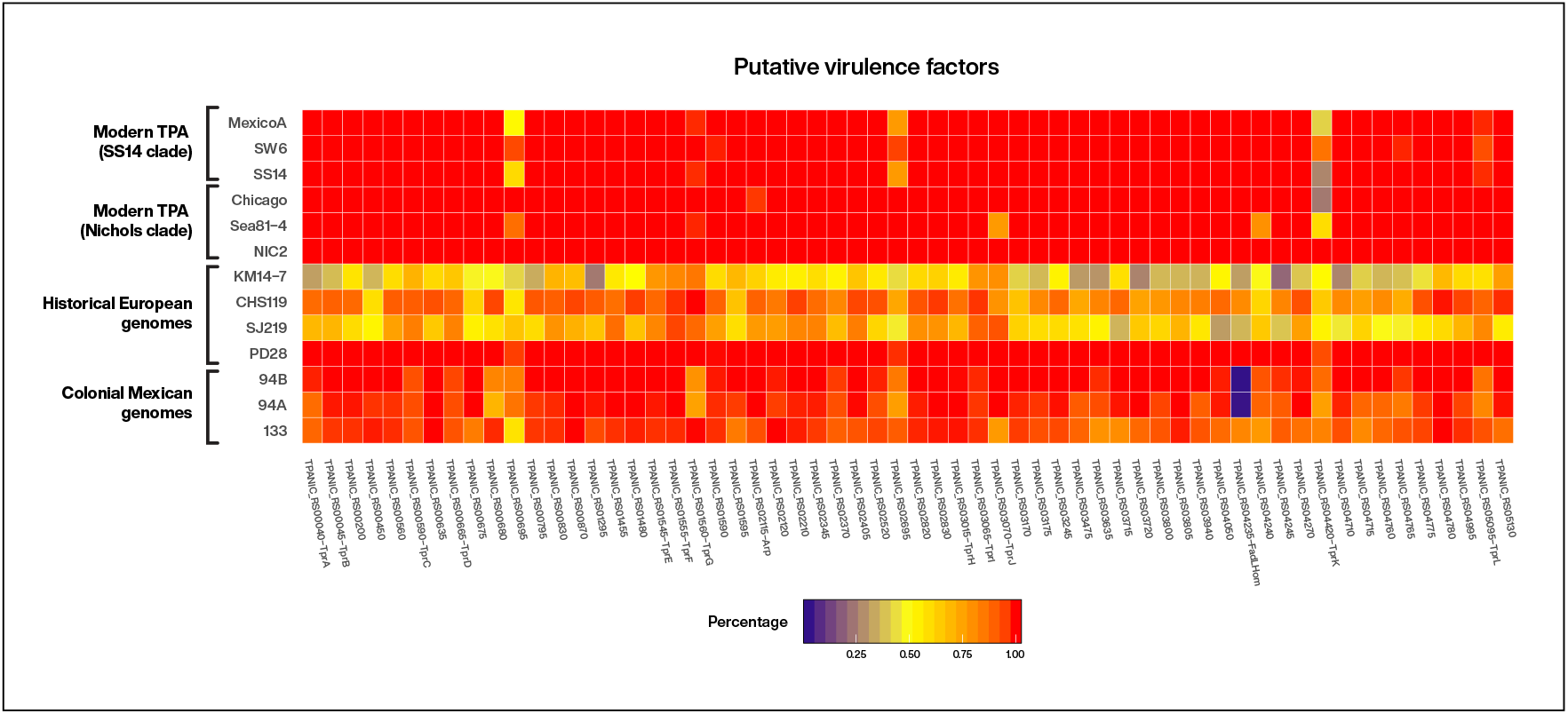
Virulence analysis of 60 putative virulence genes conducted on the four historical European genomes based on Nichols reference genome (NC_021490.2) annotation, in comparison with a selection of six modern TPA strains, three from Nichols and SS14 clades each, and the previously published colonial Mexican genomes (Schuenemann et al., 2018).

Divergence times and substitution rates were estimated using BEAST v. 2.6 ^110^. Analyses were performed under an HKY substitution model with Γ-distributed rate heterogeneity ^111,112^ and a Bayesian skyline plot demographic model (tree-prior) with 10 groups ^113^. To calibrate the clock, an uncorrelated lognormally distributed relaxed clock model ^114^ was used to allow for rate variation among lineages. An exponentially distributed prior with mean 5 × 10^−7^ s/s/y was placed on the mean clock rate. To allow for uncertainty in the sampling dates of historical genomes uniform priors across the date range defined by radiocarbon dating were placed on their ages (Supplementary table 6.B). Default priors were used for all other model components. To confirm clock-like evolution we performed a Bayesian date randomization test ^107,109,115^ (DRT) by permuting sampling dates across genomes and repeating the analysis. We performed 50 replicates and assessed significance by comparing the molecular clock rate estimates of the replicates to those estimated under the true sampling dates. As in the permutation test for the root-to-tip regression analysis above, we fixed the sampling dates of historical genomes to the middle of the date range defined by radiocarbon dating.

MCMC chains were run for 50 million steps and parameters and trees sampled every 5000 steps. Convergence was assessed in Tracer ^116^ after discarding 30% of the chains as burn-in and Treeannotator was used to compute MCC trees of the resulting posterior tree distributions. Results were visualized in R using ggplots2 ^117,118^, ggtree ^119^ and custom scripts.

### Virulence analysis

Virulence factors represented by the four ancient European genomes were assessed through a gene presence/absence analysis as described by Valtueña and colleagues ^81^. A set of 60 sequences previously associated with putative virulence functions ^45,82,83,85^ were examined based on the annotated Nichols reference genome (NC_021490.2) and without preliminary quality filtering. The coverage over each gene was calculated using genomeCoverageBed in BEDTools version 2.250 ^120^. The heatmap visualization of the gene-by-gene coverage of reads was created using ggplot2 package in R ^117,118^).

## Acknowledgments

We would like to thank Abigail Breidenstein for the proof-reading of the manuscript, and Gemeente Zwolle for the permission to sample the Kampen skeletal material. The figure graphics and layouts were designed by Michelle O’Reilly at the Max Planck Institute for the Science of Human History, Jena.

This work was supported by the Swiss National Science Foundation: grant number 188963 – “Towards the origins of syphilis” (V.J.S., K.M.), the Mäxi Foundation, Switzerland (V.J.S), the University of Zurich’s University Research Priority Program “Evolution in Action: From Genomes to Ecosystems” (V.J.S, J.N.), the Max-Planck Society (J.K.), the Senckenberg Centre for Human Evolution and Palaeoenvironment (S-HEP), University of Tübingen, Germany (V.J.S., J.K.) the Emil Aaltonen Foundation (K.M.), the Kone Foundation (K.M., P.O.) and Aatos Erkko Foundation (K.M., P.O.), Otto A. Malm Foundation (K.S.) and University of Helsinki Research Foundation (K.S.), the European Research Council under the Seventh Framework Programme (FP7/2007-2013), grant agreement 614725-PATHPHYLODYN (L.d-P.), the Oxford Martin School (L.d-P.), University of Tartu, Institute of Genomics project “Natural selection and migrations in shaping human genetic diversity in East European Plain. An ancient DNA study” (PRG243) (H.V., A.K., M.M.) and grants BFU2017-89594R from MICIN and Prometeo2016-0122 from Generalitat Valenciana (M.P-D. and F.G.-C.).

## Data Accessibility

Raw sequencing data will become available on ENA (project accession ID PRJEB35855) upon publication.

## Authors’ Contributions

V.J.S, K.M. and J.K conceived and designed the study. K.S., R.S., S.I., M.O., H.V., M.M., A.K., provided samples and archaeological context. V.J.S., F.G-C., D.K. and J.K. supervised the work. S.P., K.M., G.A. performed the experimental work. K.M., J.N., A.K., L.d-P., M.P-D., F. G-C., D.K. analyzed the sequenced data. K.M. and V.J.S. wrote the manuscript with input from all authors. All authors reviewed the manuscript.

