## Supplemental_Information_1 for "Ancient bacterial genomes reveal a formerly unknown diversity of *Treponema pallidum* strains in early modern Europe"

#### Supplementary Notes

##### Supplementary Note 1. Archaeological context and selection criteria of the samples

Kerttu Majander, Kati Salo, Sarah Inskip, Rachel Schats, Martin Malve, Aivar Kriiska

The archaeological context of Porvoo Dome cemetery, Finland, is considered to date from 17th to 18th century (K. H. Salo & Others, 2016). This time scale is confirmed by radiocarbon dating lower bound, although the upper bound reaches all the way to the modern times. It is, however, known that the cemetery was formally used until 1789, and the churchyard leveled in 1791, providing a historical limit to the remains excavated from the site (Hiekkanen, 2003; K. Salo, n.d.).

The archaeological context of Julin's plot in Turku, Finland, spans over a minimum of two hundred years, from the end of 14$^{th}$ century to mid-17$^{th}$ century (Pihlman, n.d.). The reservoir effect corrections were calculated at the Helsinki Natural History Museum Laboratory of Chronology for the individual CHS119, but these calculations reveal a dual peak pattern from 15$^{th}$ to 16$^{th}$ century and from 16$^{th}$ to 17$^{th}$ century, not securely confirming either dating (Etu-Sihvola et al., 2019; Oinonen, 2019). For detailed description of the reservoir effect correction applied, see **Supplementary note 4. Reservoir Effect Correction for the CHS119 sample**).

The Tartu context is archaeologically dated to the North-European Medieval period, and the AMS dating placing the individual in the 15$^{th}$ century confirms this estimate (Malve, 2020). Apart from the individual SJ219, a preserved fragment from her wooden coffin was dated in order to estimate a date independent from the potential dietary reservoir effect. Unfortunately, this dating provided a wider temporal range than that of the human remains from the grave, i.e. from 15$^{th}$ to 17$^{th}$ century CE.

The Kampen sample KM14-7 was radiocarbon dated to late 15th to early 17th century. The sample stems from a collection of disarticulated skeletal material found on a large excavated area within the cemetery of Gertrude's Infirmary in Kampen, the Netherlands. The use of this graveyard spans from the mid-14th to early 17th century. Reservoir effect corrections are not planned at this time. It is, however, noted from the stable isotope values ($\delta^{15}N$ and $\delta^{13}C$) in the bone material, that a marine reservoir effect due to the individual's diet could possibly affect the radiocarbon dates retrieved for the sample.

For a summary of archaeological site information and radiocarbon dating results, see **Supplementary Table 1.a) Extended sample information.**

Samples included in this study include a neonate petrous bone from Porvoo, Finland (PD28), a premolar from Turku, Finland (CHS119), a carpal bone from Tartu, Estonia (SJ219), and a tibia from Kampen, the Netherlands (KM14-7). The skeletal elements chosen were presumed the ones most likely to yield pathogen DNA, either due to the visible lesions or to the individual's perinatal death: a petrous part of the skull was used for the Porvoo neonate, as the generally most well preserved part of the skeleton. Since congenital syphilis is considered to be systemic (Ilagan et al., 1993), we expected all of the skeletal elements of this individual to be involved in the potential infection.

#### Supplementary Note 2. Radiocarbon dating and reservoir effect corrections

Kerttu Majander, Markku Oinonen, Sarah Inskip, Rachel Schats

The samples positive for treponemal DNA were radiocarbon dated in Klaus-Tschira-Archäometrie-Zentrum am Curt-Engelhorn-Zentrum, Mannheim, Germany, for all the samples, and independently in the Laboratory of Chronology, Finnish Natural History Museum, Helsinki for the sample CHS119, and the AMS laboratory, ETH Zürich for the sample SJ219. The two additional datings confirmed the initial results from Klaus-Tschira-Archäometrie-Zentrum laboratory. The results of these procedures are visualized in C14 dating curves from the OxCal program version 4.3.2 (Ramsey, 1995) as follows.

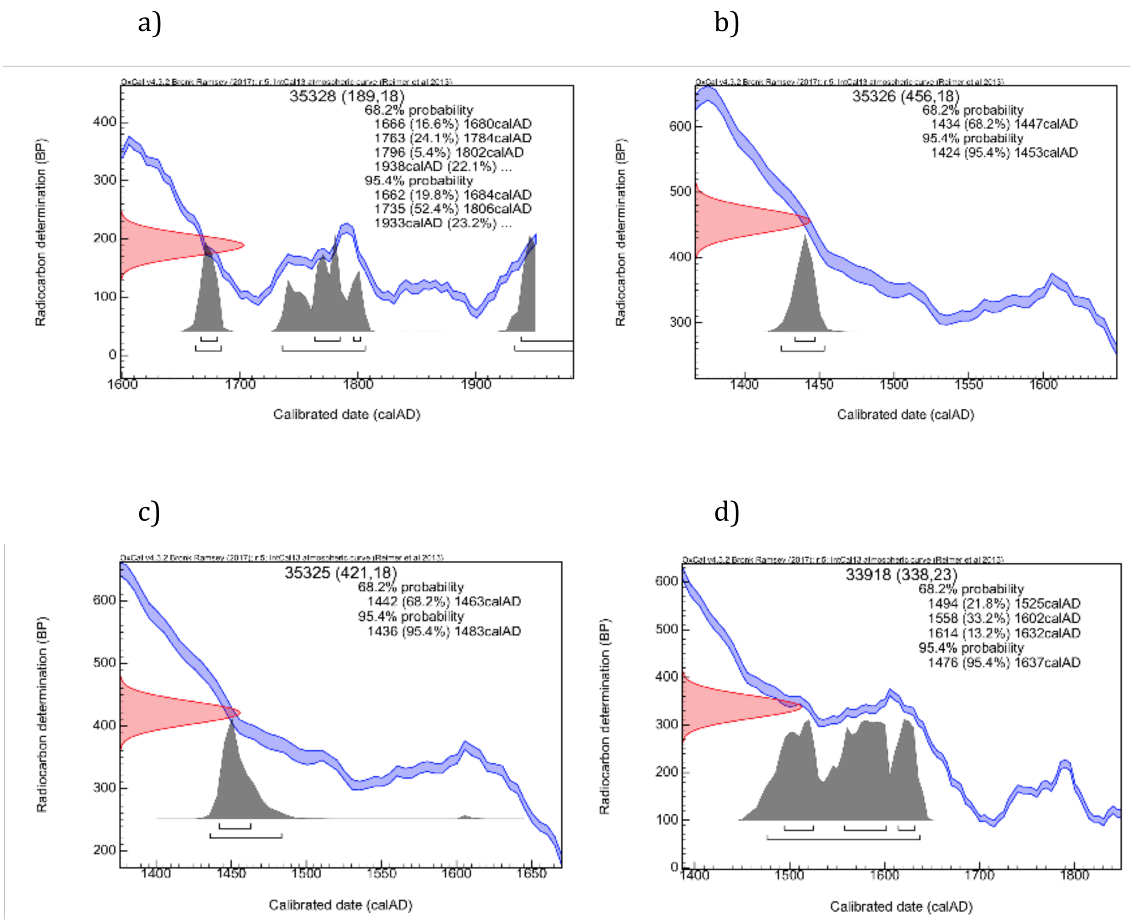


Radiocarbon curves for the samples a) PD28, b) SJ219, c) CHS119 and d) KM14-7.

#### Reservoir effect correction of the CHS119 sample

The protocols for radiocarbon dating and stable isotopic measurements for CHS119 sample at the Laboratory of Chronology, Finnish Natural History Museum, were as follows. The method for collagen extraction was based on the Longin method (Longin, 1971) and followed a previously described protocol by Berglund and colleagues (Berglund et al., 1976). To check post-mortem alteration, the contents of N and C (wt-%) and the atomic C/N ratio of the collagen were monitored (C-% = 37.4%, N-% = 13.6%, C/N ratio = 3.2) fulfilling the accepted quality criteria for well-preserved collagen i.e. 2.9–3.6 (van Klinken, 1999).

For radiocarbon analysis, the collagen sample was packed inside a silver cup (Elemental Microanalysis D2001) and the packed sample were combusted with an Elemental Analyzer (Thermo Scientific Flash 2000 NC). The resulting $CO_2$ was cryogenically trapped and reduced to graphite in the presence of zinc powder and iron catalyst (Slota et al., 1987) by using the Labview controlled HASE facility (Palonen et al., 2013). The graphite sample was measured for radiocarbon contents at the Helsinki AMS facility (Tikkanen, 2004). The result has been normalized for isotopic fractionation by using the $\delta^{13}C$ value measured with the AMS facility. Eventually, the radiocarbon date (Hela-4271) of 383 ± 24 BP was obtained.

Dietary stable isotopic (carbon and nitrogen) ratios were measured on bone collagen parallel to AMS analyses for the sample CHS119 at the Finnish Natural History Museum facility by using EA-IRMS technique. The elemental content and isotopic composition of carbon and nitrogen were measured on a NC2500 elemental analyzer coupled to a Thermo Scientific Delta V Plus isotope ratio mass spectrometer. The raw isotope data were normalised with a two-point calibration using international reference materials with known isotopic compositions (USGS-40, USGS-41). The mean measured raw $\delta^{13}C$ and $\delta^{15}N$ values, respectively, for calibration references were -26.7 and -4.7 for USGS-40, and 36.2 and 46.5 for USGS-41. Replicate analyses of a quality control reference analysed alongside the unknowns indicate a 1σ internal precision of ≤0.1 for both $\delta^{13}C$ and $\delta^{15}N$. This process resulted in dietary isotopic values of $\delta^{13}C$ = -19.7‰ and $\delta^{15}N$ = 12.3‰.

To obtain an estimate for a potential reservoir effect (RE) within the radiocarbon age, dietary modellings with FRUITS software (Fernandes et al., 2014) were performed based on an assumption of three food groups (marine, freshwater and terrestrial foods) and adopting the macronutrient concentrations, isotopic offsets and the isotopic baseline data from (Oinonen, 2019) The δIANA database (Etu-Sihvola et al., 2019) was used for gathering the Northern European isotopic data. The marine isotopic signature and corresponding marine reservoir effect was assumed to come from the vicinity of Finland Proper (Bothnian Sea, Archipelago and its surroundings). It was estimated (Oinonen, 2019) that this corresponds to the maximal marine reservoir effect (in marine animals) of 173 ± 41 $^{14}C$ years. The dietary modelling provided the contribution of bone collagen carbon from such a source and provided means to scale down the potential reservoir age of human bone collagen. Correspondingly, the maximal freshwater reservoir effect (in fish) was estimated as 107 ± 52 $^{14}C$ years. Eventually, the human bone collagen REs was estimated to be MRE = 28 ± 11 $^{14}C$ years and FRE = 11 ± 9 $^{14}C$ years yielding to a RE-corrected radiocarbon age of 345 ± 29 $^{14}C$ years corresponding to the Hela-4271 measurement. As the same amount of RE is assumed also for MAMS-35325 measurement (421 ± 18 BP) the same correction was adopted yielding to 383 ± 24 $^{14}C$ years. Finally, the combined RE-corrected radiocarbon age was obtained as 368 ± 19 $^{14}C$ years. These ages were calibrated according to Bronk Ramsey et al. (Bronk Ramsey et al., 2009) and Reimer et al (Reimer et al., 2013).

Calibrated $C^{14}$ dating curves for the sample CHS119

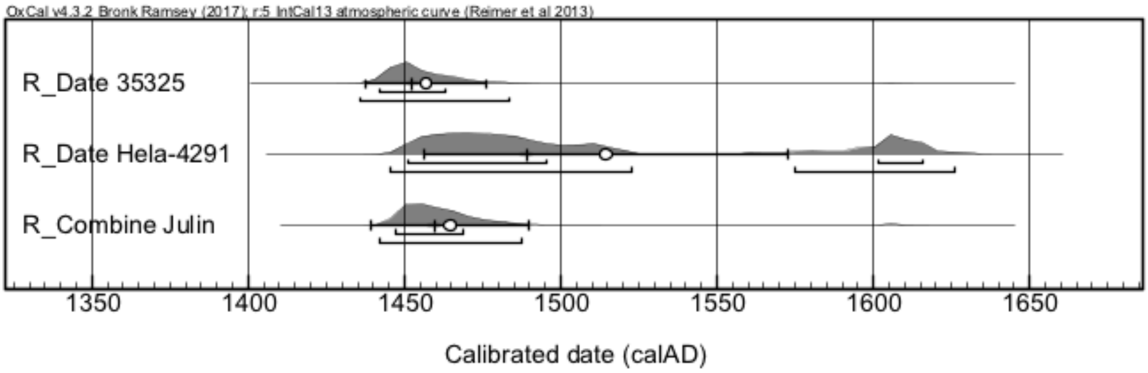


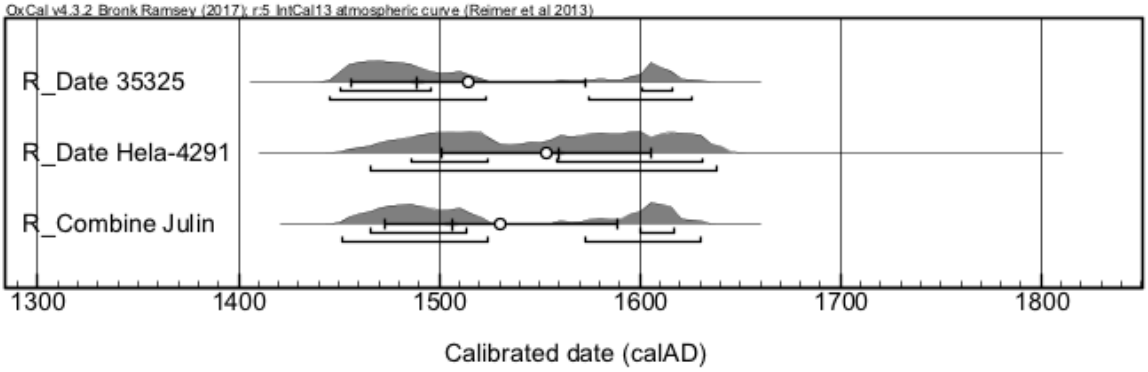


Radiocarbon dating result for the sample CHS119 prior to the reservoir effect correction

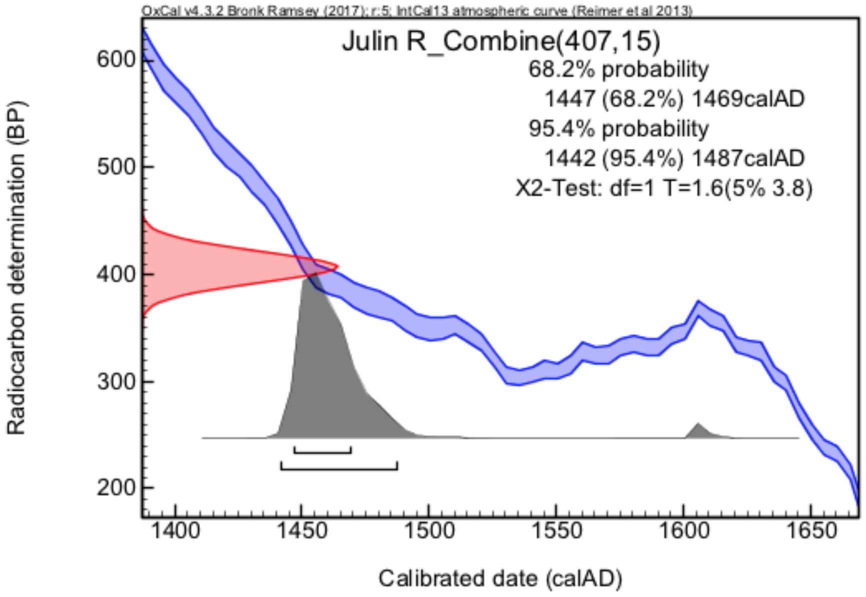

Radiocarbon dating result for the sample CHS119 with the reservoir effect correction calculated

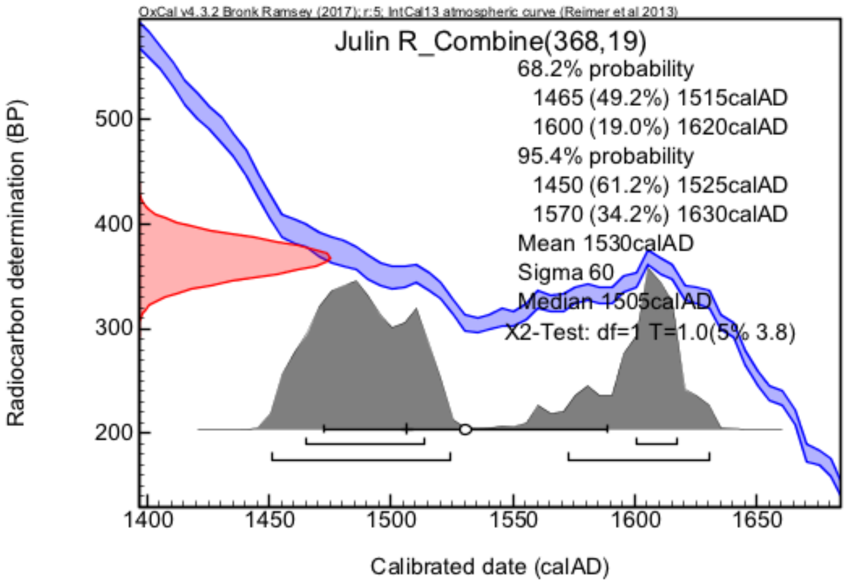


#### Supplementary Note 3. Robustness of molecular clock dating

Louis du Plessis, Arthur Kocher, Denise Kühnert, Kerttu Majander

To test the robustness of the molecular clock dating results we repeated the analyses using different combinations of demographic (constant population size coalescent, exponential growth coalescent, Bayesian skyline plot with 10 groups) and clock (strict and relaxed) models. Default priors were used for all demographic models. The same prior was used for the strict clock rate as for the mean clock rate in analyses with a relaxed clock model (exponential distribution with mean $5 \times 10^{-7}$s/s/y). Furthermore, to test the effect of constraining the sampling dates of historical genomes to the date ranges defined by radiocarbon dating (referred henceforth as narrow uniform priors) we repeated all analyses using wide uniform priors between 1000 CE and 2000 CE).

The demographic model had little effect on the estimated molecular clock rate, divergence times and sampling dates (Supplementary figures 8 and 9). Nonetheless a constant effective population size leads to slightly older divergence date estimates. The demographic models tested represent very different scenarios (constant effective population size, exponential growth and flexible growth and decline). Since all three models give similar estimates, we are confident that our results are robust to the choice of demographic model.

Estimates under the different clock models are largely overlapping. A relaxed clock model leads to wider HPD intervals than a strict clock and thus represents the more conservative choice. Furthermore, the HPD interval of the coefficient of variation (of the clock rate) estimated under a relaxed clock model does not include 0, indicating strong evidence for rate variation among lineages.

Relaxing the constraints on the sampling dates of historical genomes leads to more recent divergence time estimates more in line with previous analyses (Arora et al., 2016) and a faster clock rate estimate that is similar to the rate reported by Beale et al. (2019) for TPA (Beale et al., 2019). Similarly, the sampling date estimates of historical genomes are more recent. Nevertheless, posterior distributions of the PD28, SJ219 and 133 sampling dates place considerable weight on the date range defined by radiocarbon dating. On the other hand, the sampling date HPD interval of CHS119 falls almost entirely outside of the radiocarbon date range.

### Supplementary Figures

A)

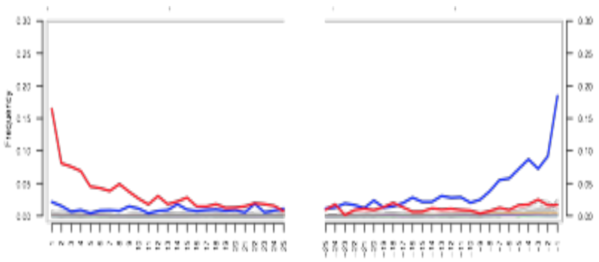

B)

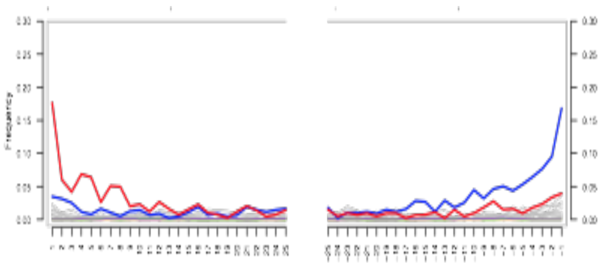

C)

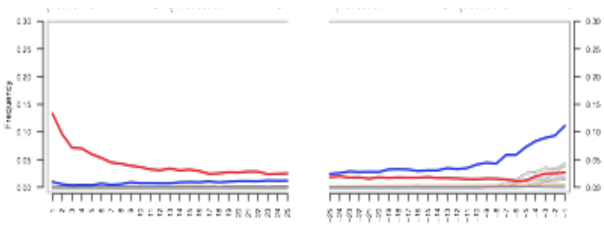

D)

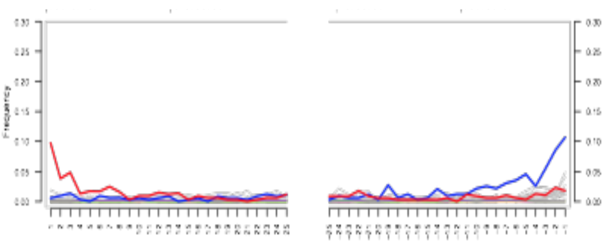

E)

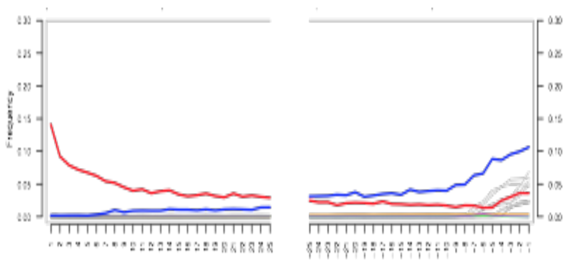

F)

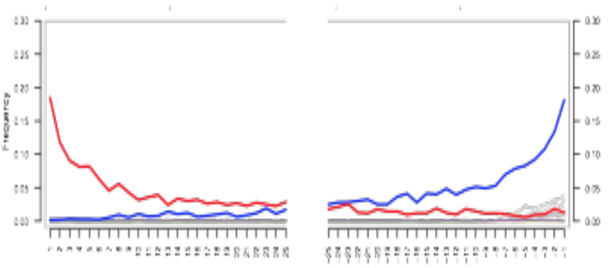

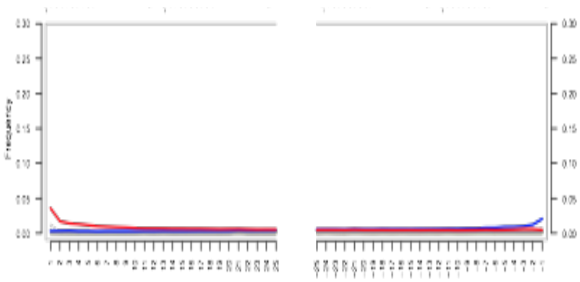

**G)**

**Supplementary Figure 1**: Misincorporation patterns from MapDamage program (Jónsson et al., 2013)  of the four *Treponema pallidum* positive samples in this study: A) mitochondrial and B) *T. pallidum* capture data for KM14-7 sample, C) mitochondrial and D) *T. pallidum* capture data for CHS119 sample, E) mitochondrial and F) *T. pallidum* capture data for SJ219 sample and G) *T. pallidum* capture data for PD28 sample. A pattern of cytosine-to-thymine base misincorporations accumulated at the end of the reads is indicative of authentic ancient DNA in the sample. The misincorporation plots show data only from libraries created without UDG treatment which is used for removal of the damaged bases. The sample PD28 shows very little damage, probably due to the relatively recent past from which the individual originates from, and the well-preserving petrous bone material used.

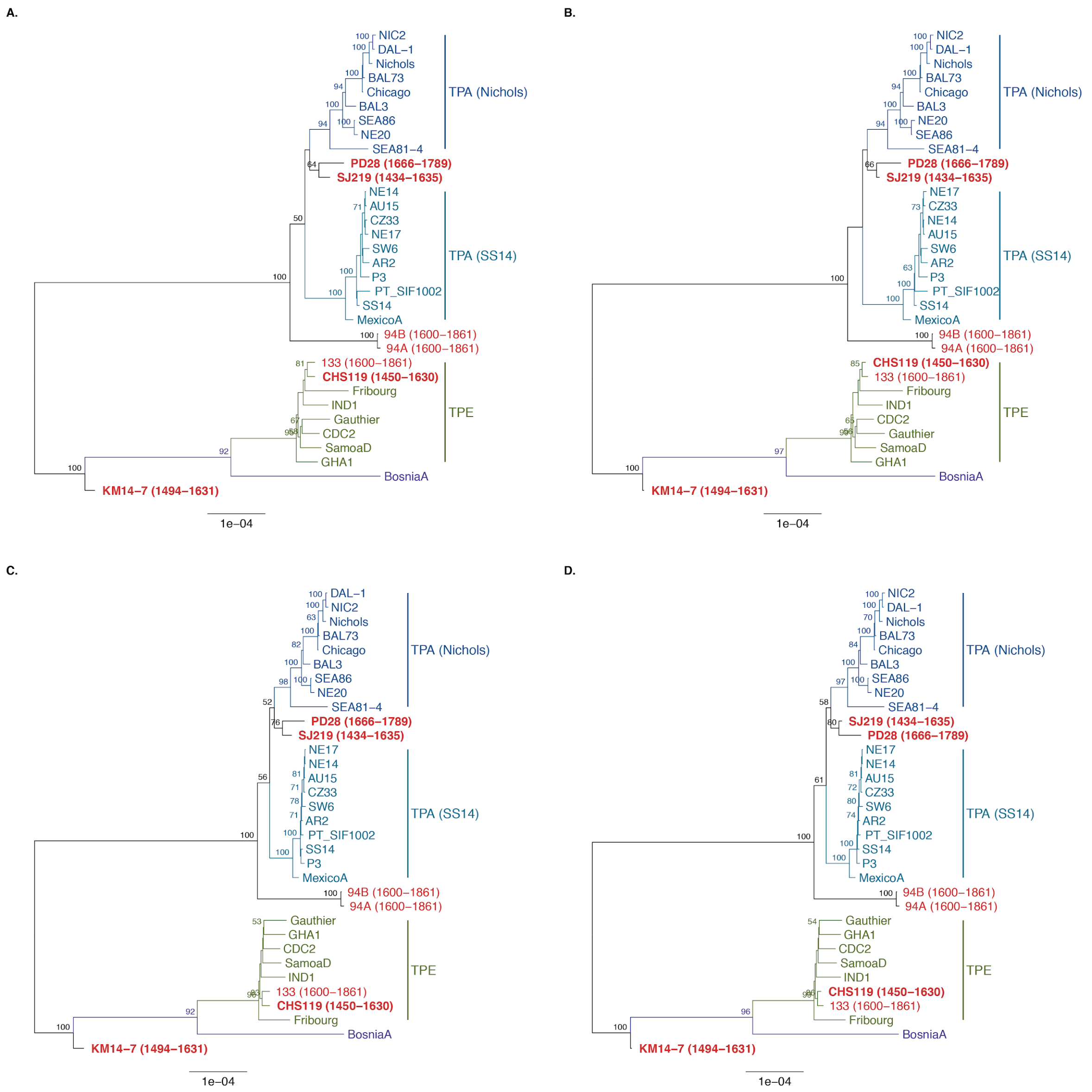


**Supplementary Figure 2:** Comparison of phylogenetic trees of *T. pallidum* based on :(A) the full alignment and no positions excluded, (B) the SNP alignment and no positions excluded, (C) the full alignment and positions with >25% missing data excluded or (D) the SNP alignment and positions with >25% missing data excluded. The trees were estimated by maximum likelihood and are midpoint-rooted. Branch lengths represent numbers of substitutions per site. Ancient genomes are marked in red and the ones generated in this study in bold text. The data derives from a set of 26 modern treponemal genomes (Arora et al., 2016; Čejková et al., 2012; Giacani et al., 2014, 2010; Pětrošová et al., 2013, 2012; Štaudová et al., 2014; Zobaníková et al., 2012; Zobanikova et al., 2013), all colonial period Mexican genomes from previously published study (Schuenemann et al., 2018) and all four historical treponemal genomes sequenced for this study.

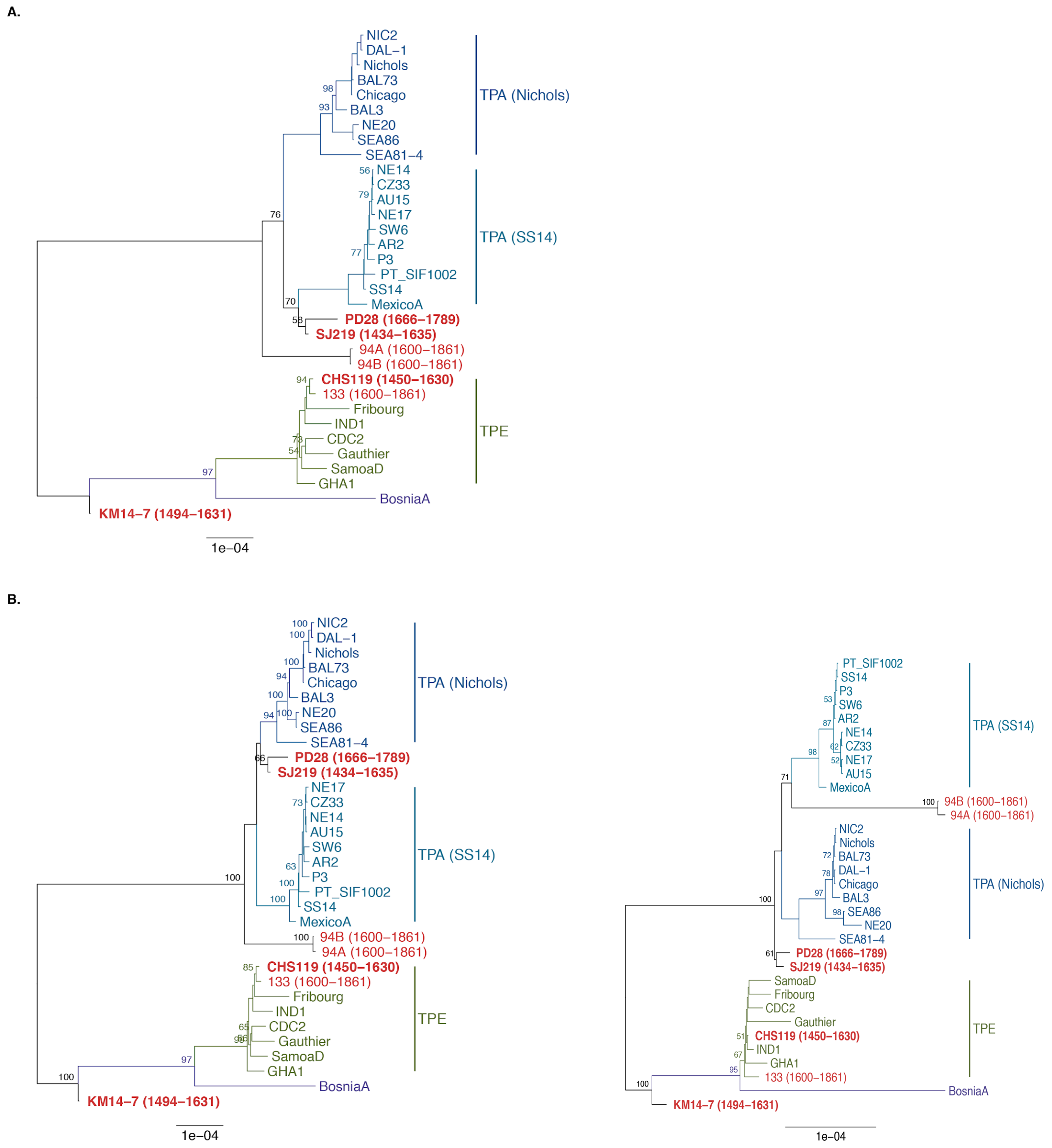


**Supplementary Figure 3**: Comparison of phylogenetic trees of *T. pallidum* based on :(A) the SNP alignment including identified recombining loci, (B) the SNP alignment with recombining loci removed. The trees were estimated by maximum likelihood and are midpoint-rooted. Branch lengths represent numbers of substitutions per site. Ancient genomes are marked in red and the ones generated in this study in bold text, (C) Phylogenetic tree based only on SNPs that were resolved in KM14-7 (141 positions). The tree was estimated by maximum likelihood and is midpoint-rooted. Branch lengths represent numbers of substitutions per site. Ancient genomes marked are in red and the ones generated in this study in bold text.

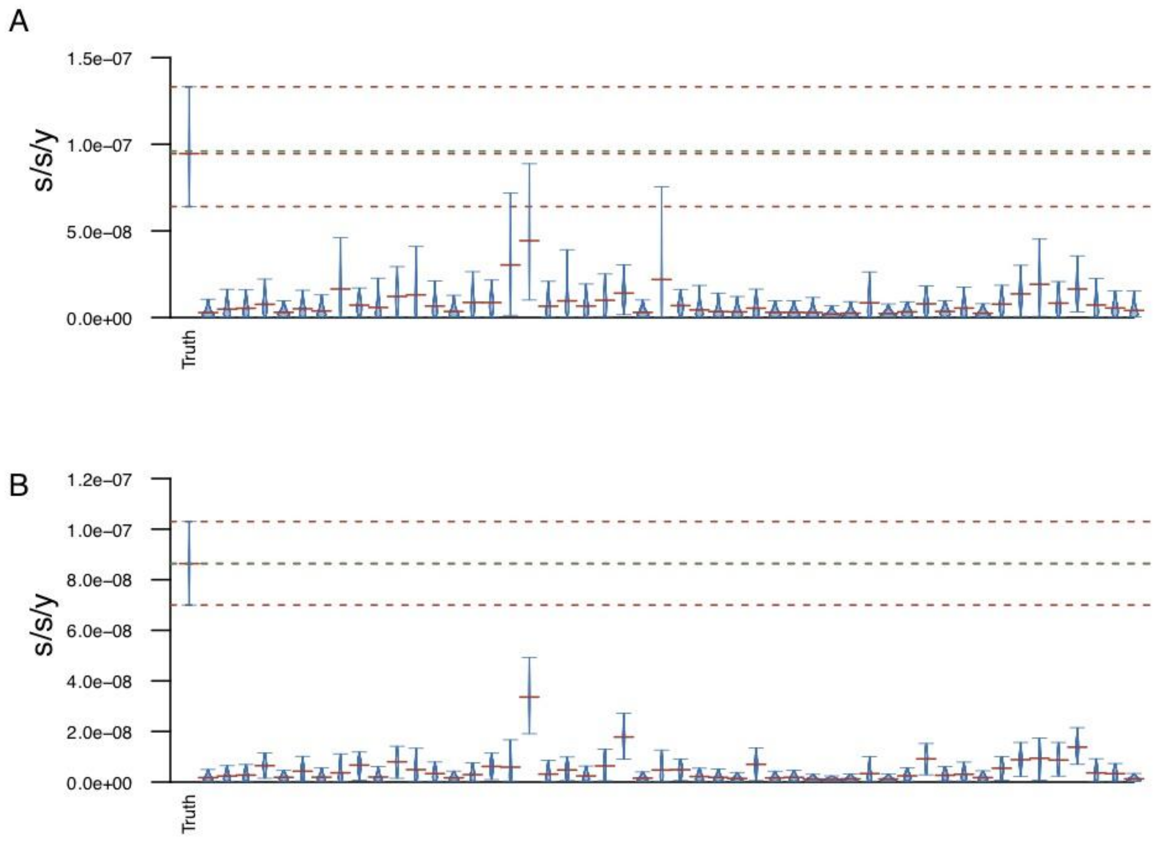


**Supplementary Figure 4:** Results of the Bayesian date randomization test (DRT) performed on the 28-genome dataset used for molecular clock dating, with 50 replicates. All analyses were performed under a Bayesian skyline plot tree prior, using either a relaxed **(A)** or a strict clock **(B)** model. The plot shows the posterior distributions for the (mean) clock rates, truncated at the upper and lower limits of the 95% HPD interval. Horizontal red lines indicate the medians of the posterior distributions. The green dashed line indicates the median (mean) estimate and red dashed lines signify the upper and lower limits of the 95% HPD interval of the clock rate inferred under the true sampling dates.

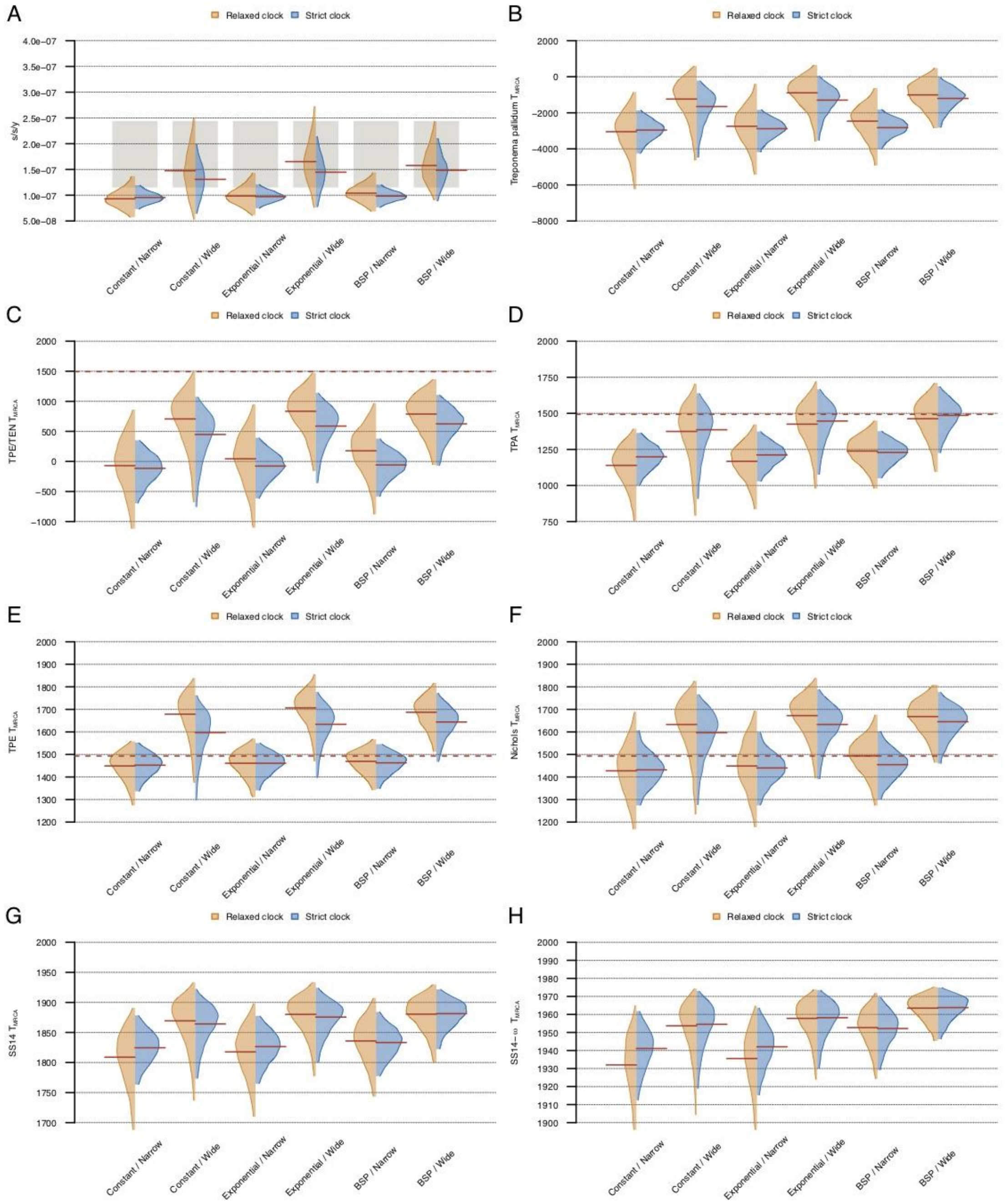


**Supplementary Figure 5:** Posterior distributions of the mean molecular clock rate **(A)** and divergence dates of clades in the tree **(B-H)** inferred under a relaxed (orange) and strict (blue) clock models with different demographic models (constant population size, exponential growth, Bayesian skyline plot) and priors on the sampling dates of historical genomes (narrow and wide uniform distributions). The distributions are truncated at the upper and lower limits of the 95% HPD interval and the red lines indicate the median estimates. The shaded boxes in **(A)** represent the substitution rate estimates reported in Beale et al. (2019) for the TPA clade. The red horizontal dashed line indicates the year 1493.

### Supplementary Tables

**Supplementary Table 1.** Extended sample information on statistical values of **a)** archaeological and radiocarbon dating analysis data, **b)** genetic data from treponemal capture **c)** genetic data mitochondrial capture.

**a)**

| Sample name (this study) | Laboratory ID | Dating laboratory ID | Grave/individual ID | Skeletal element (TP positive) | Source collection | Country of origin | C14 dating estimate (from the year CE) | C14 dating estimate (to the year CE) | Alternative dating estimate (to year CE) |
|---|---|---|---|---|---|---|---|---|---|
| PD28 | TU41 | MAMS 35328 | Grave 28 | **petrous bone** | Porvoo Dome, Porvoo | Finland | 1666 | 1950 | 1789 (archaeology) |
| SJ219 | TU578<br>TU579 | MAMS 35326 | Individual 219 | premolar,<br>**proximal phalanx** | St. Jacob's cemetery, Tartu | Estonia | 1434 | 1446 | 1635 (wood sample C14 upper bound) |
| SG34 | TU580<br>TU581 | MAMS 35327 | Individual 34 | molar,<br>metacarpale | St George's cemetery, Tartu | Estonia | 1657 | 1950 | - |
| CHS119 | TU590<br>TU591<br>TU592 | MAMS 35325 | Individual 119 | **premolar,**<br>jawbone, lower<br>jawbone, upper | Crypt of Holy Spirit, Turku | Finland | 1443 | 1460 | 1630 (aquatic reservoir effect corrected upper bound) |
| CHS101 | TU582<br>TU583 | - | Individual 101 | molar x 2 | Crypt of Holy Spirit, Turku | Finland | - | - | - |
| CHS107 | TU587<br>TU588<br>TU589 | - | Individual 107 | jawbone, upper<br>molar<br>jawbone, lower | Crypt of Holy Spirit, Turku | Finland | - | - | - |
| CHS305 | TU584<br>TU585<br>TU586 | - | Individual 305 | jawbone, upper<br>molar<br>jawbone, lower | Crypt of Holy Spirit, Turku | Finland | - | - | - |
| CHS14 | TU593 | - | Individual 14 | femur | Crypt of Holy Spirit, Turku | Finland | - | - | - |
| KM14-7 | TU391 | MAMS 33918 | Individual KM14 (disintegrated bone: 7) | **tibia** | Gertrude's Infirmary, Kampen | Netherlands | 1494 | 1631 | - |
| Cattle | Tartu A1 | ETH-100447 | Animal sample 1 | femur | Tartu | Estonia | 1268 | 1381 | - |
| Swine | Tartu A2 | ETH-100448 | Animal sample 2 | carpus | Tartu | Estonia | 1262 | 1296 | - |
| Wood | Tartu W1 | ETH-101915 | Grave 219 (fragment of coffin) | femur | Tartu | Estonia | 1463 | 1635 | - |

**b)**

| Sample name | Genome coverage X1 in % | Genome coverage X3 in % | Genome coverage X5 in % | # Raw reads | # mapped reads before duplication removal | # mapped reads after duplication removal | Endogenous DNA % | DMG 1st Base 3' (non-UDG) | DMG 2nd Base 3' (non-UDG) | DMG 1st Base 5' (non-UDG) | DMG 2nd Base 5' (non-UDG) | average fragment length | GC content in % |
|---|---|---|---|---|---|---|---|---|---|---|---|---|---|
| PD28 | 98.09 | 98.06 | 98.04 | 98204923 | 46711317 | 1430292 | 47.64 | 0.0206 | 0.0121 | 0.036 | 0.0172 | 59.91 | 52.85 |
| SJ219 | 64.31 | 15.69 | 2.58 | 193609543 | 2869982 | 29198 | 1.472 | 0.1068 | 0.0854 | 0.0974 | 0.0383 | 49.12 | 52.84 |
| CHS119 | 83.32 | 42.04 | 15.13 | 255889929 | 4491932 | 52054 | 1.746 | 0.1814 | 0.1335 | 0.1849 | 0.1179 | 54.16 | 51.52 |
| KM14-7 | 46.89 | 7.93 | 1.61 | 152268578 | 2014160 | 18034 | 1.312 | 0.168 | 0.0949 | 0.1766 | 0.0602 | 51.69 | 52.22 |

**c)**

| Sample name | average coverage on mitochondrium | # Raw reads | # mapped reads before duplication removal | # mapped reads after duplication removal | Endogenous DNA % | DMG 1st Base 3' (non-UDG) | DMG 2nd Base 3' (non-UDG) | DMG 1st Base 5' (non-UDG) | DMG 2nd Base 5' (non-UDG) | average fragment length | GC content in % | Final cont est (Schmutzi) | Final cont est low (Schmutzi) | Final cont est high (Schmutzi) | MT haplogroup | Molecular Sex |
|---|---|---|---|---|---|---|---|---|---|---|---|---|---|---|---|---|
| PD28 | - | - | - | - | - | - | - | - | - | - | - | - | - | - | - | XY |
| SJ219 | 118.9106766 | 7494443 | 384669 | 29031 | 5.316 | 0.1113 | 0.0934 | 0.1334 | 0.0959 | 67.74 | 43.31 | 0.02 | 0.01 | 0.03 | HV16 | XX |
| CHS119 | 16.24503591 | 1554965 | 11937 | 4490 | 0.787 | 0.1458 | 0.1128 | 0.1354 | 0.1102 | 59.85 | 43.42 | 0.02 | 0.01 | 0.03 | J1c2c1 | XX |
| KM14-7 | 11.02196874 | 261772 | 4233 | 3414 | 1.7 | 0.1846 | 0.0916 | 0.1647 | 0.0808 | 53.35 | 44.04 | 0.01 | 0 | 0.02 | U2e1f1 | XY |

**Supplementary Table 2:** The complete dataset of modern *Treponema pallidum* strains (Arora et al., 2016; Čejková et al., 2012; Giacani et al., 2014, 2010; Pětrošová et al., 2013, 2012; Štaudová et al., 2014; Sun et al., 2016; Zobaníková et al., 2012; Zobanikova et al., 2013) and ancient genomes used in this study. The two modern strains (NIC2, Nichols), the two ancient strains from colonial Mexico(Schuenemann et al., 2018)  (94A and 94B) along with the European ancient genome KM14-7 from this study, all excluded from the final BEAST analyses, are marked with an asterisk.

| Name | Public_Accession_or_WSI-lane | type | Isolation_year | Dating estimate (CE) | geo_location | geo_country | Reference | Obtained from |
|---|---|---|---|---|---|---|---|---|
| NIC2* | SRR3268713 | public | 1912 | | NA | USA | Arora 2016 | raw sequencing data |
| DAL_1 | NC_016844.1 | public | 1991 | | Dallas | USA | Zobaníková 2012 | raw sequencing data |
| Nichols* | NC_021490.2 | public | 1912 | | NA | USA | Pětrošová 2013 | raw sequencing data |
| BAL73 | SRR3268726 | public | 1973 | | Baltimore | USA | Arora 2016 | raw sequencing data |
| Chicago | NC_017268.1 | public | 1951 | | Chicago | USA | Giacani 2010 | GenBank |
| BAL3 | SRR3268715 | public | 1973 | | Baltimore | USA | Arora 2016 | raw sequencing data |
| SEA86 | SRR3268740 | public | 1986 | | Seattle | USA | Arora 2016 | raw sequencing data |
| NE20 | SRR3268710 | public | 2013 | | NA | Netherlands | Arora 2016 | raw sequencing data |
| SEA81_4 | 15169_7#74 | public | 1981 | | Seattle | USA | Giacani 2014 | GenBank |
| AU15 | SRR3268722 | public | 2013 | | NA | Austria | Arora 2016 | raw sequencing data |
| NE14 | SRR3268703 | public | 2013 | | NA | Netherlands | Arora 2016 | raw sequencing data |
| CZ33 | SRR3268690 | public | 2013 | | NA | Czech Republic | Arora 2016 | raw sequencing data |
| NE17 | SRR3268707 | public | 2013 | | NA | Netherlands | Arora 2016 | raw sequencing data |
| AR2 | SRR3268682 | public | 2013 | | NA | Argentina | Arora 2016 | raw sequencing data |
| SW6 | SRR3268732 | public | 2012 | | NA | Switzerland | Arora 2016 | raw sequencing data |
| P3 | SRR2996731 | public | 2016 | | NA | China | Sun 2016 | GenBank |
| PT_SIF1002 | NZ_CP016051.1 | public | 2011 | | NA | Portugal | Pinto 2016 | GenBank |
| SS14 | NC_021508.1 | public | 1977 | | Atlanta | USA | Pětrošová 2013 | GenBank |
| MexicoA | NC_018722.1 | public | 1953 | | NA | Mexico | Pětrošová 2012 | GenBank |
| Fribourg | NC_021179.1 | public | 1966 | | NA | Guinea | Zobaníková | GenBank |
| IND1 | SRR3268698 | public | 1990 | | NA | Indonesia | Arora 2016 | raw sequencing data |
| Gauthier | NC_016843.1 | public | 1960 | | NA | Ghana | Čejková 2012 | GenBank |
| CDC2 | NC_016848.1 | public | 1980 | | Akorabo | Ghana | Čejková 2012 | GenBank |
| SamoaD | NC_016842.1 | public | 1953 | | NA | Western Samoa | Čejková 2012 | raw sequencing data |
| GHA1 | Ghana-051 | public | 1988 | | NA | Ghana | Arora 2016 | raw sequencing data |
| BosniaA | SRR3268694 | public | 1950 | | NA | Bosnia | Štaudová 2014 | GenBank |
| 133 | PRJEB21276 | public | | 1600-1861 | Mexico City | Mexico | Schuenemann 2018 | raw sequencing data |
| 94A* | PRJEB21276 | public | | 1600-1862 | Mexico City | Mexico | Schuenemann 2019 | raw sequencing data |
| 94B* | PRJEB21276 | public | | 1600-1863 | Mexico City | Mexico | Schuenemann 2020 | raw sequencing data |
| PD28 | PRJEB35855 | new | | 1666-1789 | Porvoo | Finland | this study | raw sequencing data |
| SJ219 | PRJEB35855 | new | | 1429-1635 | Tartu | Estonia | this study | raw sequencing data |
| CSH119 | PRJEB35855 | new | | 1450-1630 | Turku | Finland | this study | raw sequencing data |
| KM14-7* | PRJEB35855 | new | | 1494-1631 | Kampen | Netherlands | this study | raw sequencing data |

**Supplementary Table 3:** Results of the SNP evaluation procedure for the ancient genomes included in the study.

| Sample | Nb. SNPs called | Nb. SNPs filtered | % SNPs filtered |
|---|---|---|---|
| 133 | 607 | 17 | 2.8 |
| 94A | 294 | 20 | 6.8 |
| 94B | 368 | 3 | 0.8 |
| KM14-7 | 54 | 22 | 40.7 |
| PD28 | 244 | 0 | 0 |
| SJ219 | 70 | 18 | 25.7 |
| CHS119 | 377 | 17 | 4.5 |

**Supplementary Table 4:** Investigation of SNPs positions resolved in KM14-7 that were characteristic of TPA or TPE/TEN (for which the majority variant was differing between both clades, with a frequency >90% within each clade). For each genome, we report the number and proportion of resolved SNPs characteristic of each clade.

| name | clade | Nb. pos. resolved | TPA like (%) | TPE/TEN like (%) |
|---|---|---|---|---|
| KM14-7 | ancient | 30 | 40 | 60 |
| AR2 | TPA | 30 | 100 | 0 |
| Nichols | TPA | 30 | 100 | 0 |
| NIC2 | TPA | 30 | 100 | 0 |
| BAL3 | TPA | 28 | 100 | 0 |
| BAL73 | TPA | 29 | 100 | 0 |
| SEA81-4 | TPA | 28 | 89.3 | 10.7 |
| SEA86 | TPA | 22 | 100 | 0 |
| NE20 | TPA | 30 | 100 | 0 |
| Chicago | TPA | 30 | 100 | 0 |
| DAL-1 | TPA | 30 | 100 | 0 |
| SS14 | TPA | 30 | 100 | 0 |
| MexicoA | TPA | 30 | 100 | 0 |
| SW6 | TPA | 30 | 100 | 0 |
| AU15 | TPA | 30 | 100 | 0 |
| CZ33 | TPA | 25 | 100 | 0 |
| NE14 | TPA | 30 | 100 | 0 |
| NE17 | TPA | 30 | 100 | 0 |
| P3 | TPA | 30 | 100 | 0 |
| PT_SIF1002 | TPA | 30 | 100 | 0 |
| Gauthier | TPE/TEN | 30 | 0 | 100 |
| CDC2 | TPE/TEN | 30 | 0 | 100 |
| SamoaD | TPE/TEN | 30 | 0 | 100 |
| IND1 | TPE/TEN | 30 | 0 | 100 |
| GHA1 | TPE/TEN | 29 | 0 | 100 |
| Fribourg | TPE/TEN | 30 | 0 | 100 |
| BosniaA | TPE/TEN | 30 | 0 | 100 |
| SJ219 | ancient | 7 | 100 | 0 |
| PD28 | ancient | 30 | 100 | 0 |
| CHS119 | ancient | 17 | 0 | 100 |
| 133 | ancient | 23 | 0 | 100 |
| 94A | ancient | 25 | 84 | 16 |
| 94B | ancient | 28 | 85.7 | 14.3 |

**Supplementary Table 5.** Recombination events detected across the complete dataset of 26 modern *Treponema pallidum* strains and six ancient genomes (94A, 94B, 133 from Schuenemann *et al.* 2018 and genomes PD28, CHS119 and SJ219 from this study). Slashes are used to separate the different potential gene donor strains. The affected recipient strains are separated by commas. Arrows point to the likely direction of recombination between the donor and recipient strains. An interrogation mark indicates and uncertain, yet likely involvement in the event.

| Gene ID | Event | Ini | End | Minimal size | Strains involved |
|---|---|---|---|---|---|
| TPANIC_0136 | 3 | 329 | 394 | 65 | Yaws -> Nichols clade |
| | 4 | 404 | 422 | 18 | Yaws/Bosnia -> Nichols clade |
| | 5 | 973 | 1034 | 61 | Yaws/Bosnia/133/CHS119? -> PD28, Nichols clade |
| | 6 | 1370 | 1381 | 11 | Yaws/Bosnia/133 -> Nichols clade |
| TPANIC_0164 | 1 | 228 | 341 | 113 | Yaws/Bosnia -> Sea86, Ne20, 94A, 94B |
| TPANIC_0179 | 1 | 259 | 647 | 388 | Yaws/Bosnia/133 -> PD28, Nichols clade |
| TPANIC_0326 | 1 | 425 | 2522 | 2097 | Bosnia -> SS14 clade |
| TPANIC_0462 | 1 | 151 | 984 | 833 | Yaws/Bosnia -> Sea86, Ne20 |
| TPANIC_0488 | 1 | 523 | 1162 | 639 | Bosnia -> MexicoA |
| TPANIC_0515 | 1 | 949 | 2845 | 1896 | Yaws/Bosnia/133 -> 94A, 94B, Nichols clade |
| TPANIC_0548 | 1 | 294 | 946 | 652 | Yaws/Bosnia -> Nichols clade |
| TPANIC_0558 | 1 | 414 | 834 | 420 | Yaws/Bosnia/CHS119?/133? -> PD28, 94B, 94A?, SS14 clade, MexicoA |
| TPANIC_0865 | 1 | 258 | 576 | 318 | Yaws/Bosnia/CHS119?/133? -> Sea86, Ne20, Sea81-4 |
| | 2 | 864 | 1332 | 468 | Bosnia -> Sea86, Ne20, Sea81-4 |
| TPANIC_0967 | 1 | 429 | 1538 | 1109 | Yaws/Bosnia -> Sea81-4 |
| TPANIC_0968 | 1 | 36 | 1239 | 1203 | Bosnia -> Sea81-4 |

**Supplementary Table 6** (A) Date ranges (defined by radiocarbon dating) used for historical sequences in the molecular clock dating analyses. The mean sampling date was used in robustness analyses to test the strength of the molecular clock signal. (B) Posterior $T_{MRCA}$ estimates of clades in the 28 -genome dataset used for molecular clock dating. The posterior probability that a clade is monophyletic is calculated as the proportion of posterior trees where the clade is monophyletic. (C) Posterior sampling date estimates for historical genomes in the 28-genome dataset used for molecular clock dating. The posterior probability that a sample is pre-Columbian is calculated as the proportion of posterior samples with a date < 1493.

A)

| Name | Sampling location | Strain | Radiocarbon date range | Mean sampling date |
|---|---|---|---|---|
| 133 | Mexico | TPE | 1600–1861 | 1730.5 |
| PD28 | Finland | TPA | 1666–1789 | 1727.5 |
| SJ219 | Estonia | TPA | 1434–1635 | 1534.5 |
| CHS119 | Finland | TPE | 1450–1630 | 1540.0 |

B)

| Clade | Median | HPD lower | HPD upper | Monophyletic |
|---|---|---|---|---|
| SS14-w | 1952.70 | 1924.27 | 1971.86 | 0.98 |
| SS14 | 1835.85 | 1743.43 | 1907.45 | 0.99 |
| Nichols | 1493.48 | 1273.71 | 1676.09 | 0.98 |
| TPA | 1238.77 | 979.20 | 1449.26 | 0.98 |
| TPE | 1469.42 | 1341.95 | 1567.15 | 0.97 |
| TPE/TEN | 176.33 | -880.64 | 969.85 | 0.97 |

C)

| Sample | Median | HPD lower | HPD upper | Pre-Columbian |
|---|---|---|---|---|
| 133 | 1664.33 | 1600.04 | 1773.13 | 0.00 |
| PD28 | 1744.17 | 1678.08 | 1788.99 | 0.00 |
| SJ219 | 1536.25 | 1436.47 | 1623.41 | 0.26 |
| CHS119 | 1602.08 | 1526.41 | 1630.00 | 0.02 |
